# Odor representations are structured by heterogeneous tuning and concentration-dependence of functionally distinct olfactory sensory neuron subpopulations

**DOI:** 10.64898/2026.10.27.762248

**Authors:** Jason B. Castro, Shawn D. Burton, Joel Hallkaj, Matt Wachowiak

**Author notes:** **Correspondence:** Matt Wachowiak. the first two authors contributed equally.

## Abstract

Olfactory stimuli (odorants) are represented by combinatorial patterns of olfactory sensory neuron (OSN) activation that are translated into spatially-organized glomerular maps at their input to the CNS. How glomerular maps relate to odorant features - including chemical structure and concentration - remains a key question in understanding the neural mechanisms of odor coding. Here, we compared glomerular response maps for the same 59 odorants across three concentration regimes, using GCaMP6 imaging from OSN axon terminals in the olfactory bulb (OB) of the anesthetized mouse. At the lowest regime, in which concentrations were chosen to distinguish the most sensitive glomeruli for each odorant, glomeruli were exceptionally narrowly tuned and responses across glomeruli were extremely sparse. We then compared responses at two higher-concentration regimes, ‘low’ and ‘high’, differing by 25-fold from each other and 2 - 5 log units higher than the ‘most-sensitive’ regime. Increasing concentration broadened tuning of OSN inputs; this broadening was heterogeneous across chemical space and was well-described by classifying glomeruli into several subpopulations with distinct consensus tuning profiles. Each subpopulation was selectively tuned to one or a few odorant chemical classes and spatially clustered in a manner consistent with the projection patterns of receptor-defined OSN classes. Glomerular subpopulations retained distinct tuning and spatial localization even at the highest concentrations. OSN subpopulations thus sample chemical space nonuniformly across a large concentration range. These results suggest a lower-dimensional structure in the representation of odorant information that largely reflects OSN subpopulations which, in turn, map to olfactory receptor subfamilies. This organization may enable distinct odor coding strategies for different OSN classes and for parts of chemical feature space with distinct ethological relevance.

## Introduction

At the first synaptic level of the olfactory pathway, odorants are represented by combinatorial patterns of olfactory sensory neuron (OSN) activation. Understanding how the repertoire of OSNs represents olfactory stimulus space remains a difficult problem due to the high-dimensional nature of both olfactory stimulus space and neural coding space, where, in rodents, nearly 1000 coding ‘channels’ defined by their cognate odorant receptors (ORs) represent tens of thousands of odorous compounds varying in chemical features. OSNs expressing the same receptor converge onto the same glomerulus, and subpopulations of OSNs expressing evolutionarily-related receptor converge to nearby glomeruli (Bozza et al., 2009; Wang et al., 2022; Zhu et al., 2022). Thus, odorant-evoked patterns OSN activation are translated into spatially-organized glomerular maps at the level of input to the CNS.

Characterizing how glomerular maps relate to olfactory stimulus features can yield insights into the nature of odorant - receptor binding, population-level odor coding strategies by OSNs, and the potential organization of circuits underlying the initial processing of incoming olfactory information (Murthy, 2011). Previous studies have yielded some consensus about how odorants are represented at this level. For example, nearby glomeruli tend to have similar tuning profiles (Takahashi et al., 2004; Johnson and Leon, 2007; Ma et al., 2012) and increasing odorant concentration broadens the tuning of OSNs (Sato et al., 1994; Duchamp-Viret et al., 2000; Kajiya et al., 2001) and recruits input to more - and more highly overlapping - glomeruli (Wachowiak and Cohen, 2001; Fried et al., 2002; Hu et al., 2020). Key questions remain unresolved, however, including whether the high-dimensional space of olfactory stimuli is reduced to a lower-dimensional representation by OSNs, what the determinants of any lower-dimensional structure may be, and how such a structured representation may change with odorant concentration, which can vary widely in natural settings (Wachowiak et al., 2025). Addressing these questions has been limited in part by study-specific differences in the composition of odorant stimulus panels and the range of concentrations used.

Recently we characterized glomerular responses to 185 odorants presented at concentrations distinguishing the most-sensitive glomeruli for each odorant; these concentrations were substantially lower than those used in prior studies (Burton et al., 2022). In this regime glomeruli are exceptionally narrowly tuned, resulting in a high-dimensional but nonetheless sparsely structured representation of olfactory stimulus space by the OSN population. These results are qualitatively distinct from those of prior studies characterizing OSN and glomerular odorant representations using higher concentrations (Wachowiak and Cohen, 2001; Soucy et al., 2009; Ma et al., 2012).

To resolve these differences and gain better insight into how OSNs map olfactory stimuli to glomerular patterns across chemical feature space and odorant concentration, here we sought to compare glomerular response maps for the same set of diverse odorants across concentrations. We imaged OSN response maps across dorsal glomeruli to 59 odorants (a subset of the original 185), with each odorant presented at two concentrations differing by 25-fold. These concentrations were 100 - 10,000-fold greater than those used in our earlier study, collectively allowing us to compare odorant representations for the same 59 odorants across a total concentration range of ∼ 3 - 5 orders of magnitude.

We found that increasing odorant concentrations broadened the tuning of OSN inputs to glomeruli, as expected, but that such broadening was both non-random and heterogeneous with respect to chemical space, and was well-described by classifying glomeruli into a few subpopulations defined by their tuning in the low-concentration regime. These results point to the emergence of a low-dimensional structure in the representational space of olfactory stimuli by the OSN repertoire as odorant concentrations increase. Our analysis further suggests that this structure largely reflects OSN subpopulations defined by receptor subfamilies, which maintain a nonuniform sampling of chemical space even at the highest concentrations. This heterogeneous representation of olfactory stimuli may allow the olfactory system to differentially process chemical signals with distinct ethological relevance across a dynamic sensory environment.

## Results

### Increasing concentration heterogeneously broadens glomerular representations of odorants

To characterize how OSN tuning and glomerular representations of olfactory stimuli change as a function of odorant concentration, we used GCaMP6f expressed selectively in OSNs (OMP-Cre x RCL-GCaMP6f mice) to image odorant-evoked Ca2+ signals in OSN axon terminals in OB glomeruli. We obtained odorant response maps from the dorsal OBs of anesthetized mice using epifluorescence imaging, artificial inhalation and high-throughput olfactometry (n = 3 mice, 4 OBs), as described previously (Burton et al., 2022). We used a panel of 59 odorants, each presented at two concentrations differing by 25-fold; the 59 odorants were a subset of the 185 used previously to map glomerular tuning and spanned a similar range of chemical space (**Fig. 1A**). In the present study, we used concentrations 100 – 1000-fold higher than those used previously (**Fig. 1B**). In some cases, we also included for analysis data from (Burton et al., 2022) (shared at https://github.com/WachowiakLab/OdorMappingData), allowing us to characterize glomerular responses across 3 - 5 log units of concentration for the same 59 odorants (**Table S1**). For clarity, we refer to the two higher concentration levels tested presently as ‘low’ and ‘high’, and the lower concentrations tested in (Burton et al., 2022) as ‘most-sensitive’.

**Figure 1.**
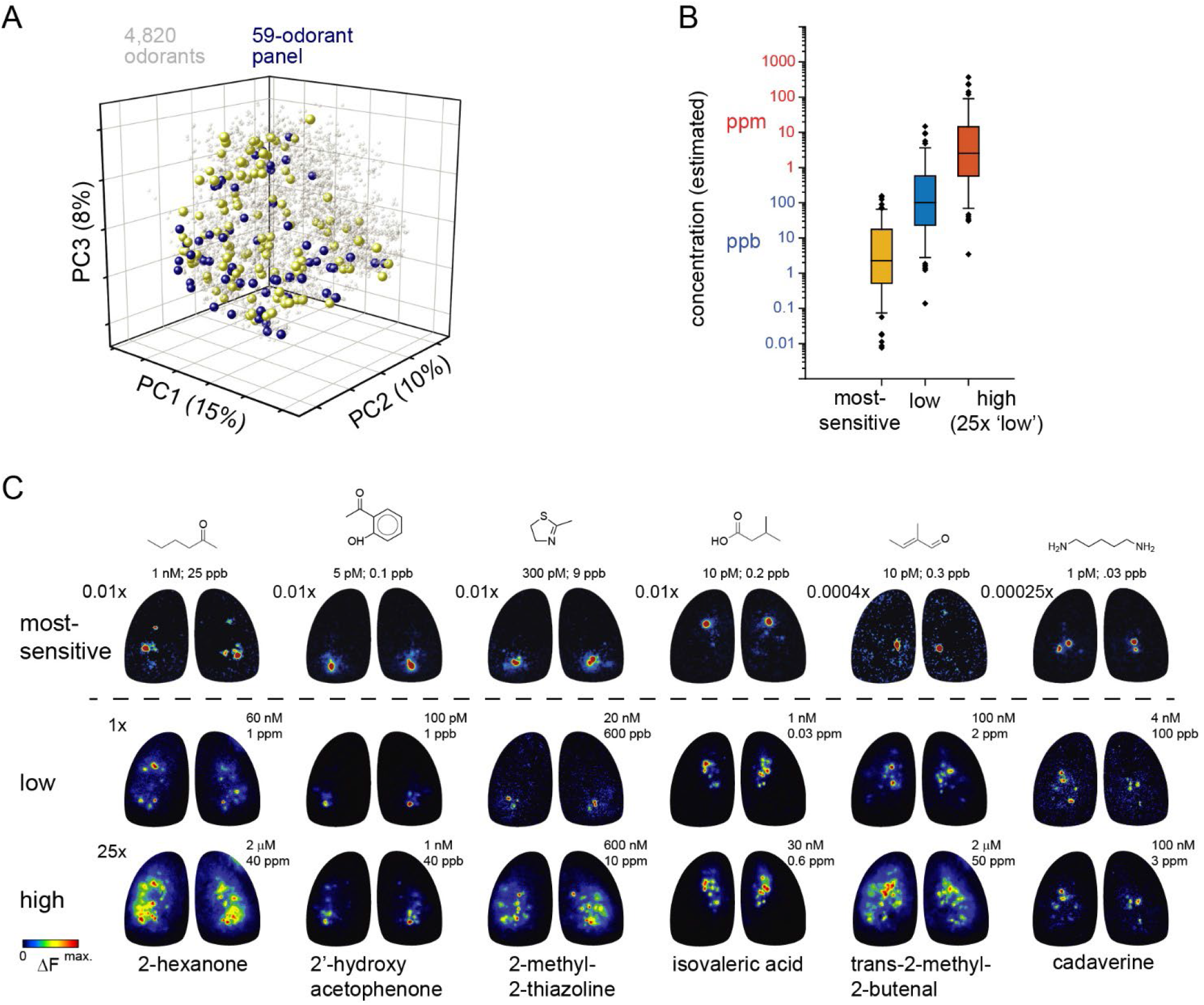
Characterizing glomerular response maps across concentration. **A.** Coverage of physicochemical feature space by the odorant panel. Points are odorant compounds projected onto the first 3 principal components (PC) of the feature space computed from the MACCS fingerprints of 4820 compounds (see Methods). The 59-odorant panel is shown in purple; the remaining 126 odorants from the previous dataset (i.e., ‘most-sensitive’) (Burton et al., 2022) are shown in yellow. Gray points indicate the remaining compounds from the OpenPOM-GSLF dataset (see Methods). **B.** Distribution of estimated delivered concentrations for the odorant panel in the three concentration regimes ‘most-sensitive’, ‘low’ and ‘high’. Box plots show median and quartiles; whiskers indicate 10 - 90th percentiles, with outliers show as individual points. **C.** Representative response maps (ΔF) evoked by single odorants presented at each of the three concentration regimes. Each column is the same odorant. Top row: ‘Most-sensitive’ concentrations, taken from (Burton et al., 2022). Concentration relative to the ‘low’ concentration is shown at the upper left of each map. Absolute delivered concentration (estimated) is shown above each map. Middle, Bottom rows: ‘Low’ and ‘High’ concentration responses, all imaged in the same mouse (but separate from ‘Most-Sensitive’ maps). Each response map is scaled from 0 to its own maximum.

Example response maps for five odorants presented at these three concentration levels are shown in **Fig. 1C**. Several qualitative trends are apparent in these examples. First, increasing odorant concentration recruits input to increasing numbers of glomeruli. Amplitudes of glomerular responses also increased with increasing concentrations (not shown), consistent with previous results (Wachowiak and Cohen, 2001; Oka et al., 2006; Storace and Cohen, 2017). Second, some odorants recruit more glomeruli, and across a larger area of the dorsal OB, than others (compare isovaleric acid vs. trans-2-methyl-2-butenal, **Fig. 1C**). Third, some degree of spatial heterogeneity in the locations of activated glomeruli persists even at the highest odorant concentrations. Together these observations suggest that the impact of odorant concentration on OSN tuning and glomerular odorant representations may vary across the OSN repertoire. We investigated the nature of these concentration-dependent changes using a variety of analyses, described in the sections below, focusing primarily on understanding concentration-dependent changes in response selectivity (i.e., tuning in chemical feature space).

We earlier reported that, at the ‘most-sensitive’ odorant concentrations, which were in the single-to sub-parts per billion (ppb) range (sub-nanomolar to picomolar; **Fig. 1B**), OSN inputs to individual glomeruli are narrowly tuned across chemical space and glomerular representations were extremely sparse (Burton et al., 2022). Consistent with previous work, increasing odorant concentration lowered the sparseness of glomerular odorant representations (Kajiya et al., 2001; Wachowiak and Cohen, 2001; Ma et al., 2012) and broadened the tuning of individual glomeruli (**Fig. 2A,B**). Population sparseness, reflecting the selectivity of activation across the glomerular population by a given odorant (see Methods), decreased from 0.99 ± 0.008 (mean ± st. dev.; n = 57 odorants, averaged by odorant across 8 OBs) for the ‘most-sensitive’ dataset to 0.97 ± 0.02 and 0.90 ± 0.06 (59 odorants, averaged by odorant across 4 OBs), respectively, for the ‘low’ and ‘high’ concentrations. The number of glomeruli activated to any degree by a given odorant increased from 2.6 ± 1.5 (median ± IQR; n = 59 odorants averaged across OBs) at the ‘most-sensitive’ concentrations to 5.75 ± 4.75 at ‘low’ concentration to 16.25 ± 10.5 at the ‘high’ concentration, corresponding to a range of ∼2 to 10% (11 ± 7%; median ± IQR) of the imaged glomerular population (**Fig. 2C**). Lifetime sparseness (S_L_), a measure of stimulus selectivity across odorants, ranging from 0 (no stimulus selectivity) to 1 (response to only a single stimulus), decreased from 0.98 ± 0.01 (n = 587 glomeruli pooled across 8 OBs) in the ‘most-sensitive’ dataset to 0.96 ± 0.04 (n = 376, 4 OBs) at the ‘low’ concentrations to 0.89 ± 0.1 (n = 481, 4 OBs) at the ‘high’ concentrations (**Fig. 2D**).

**Figure 2.**
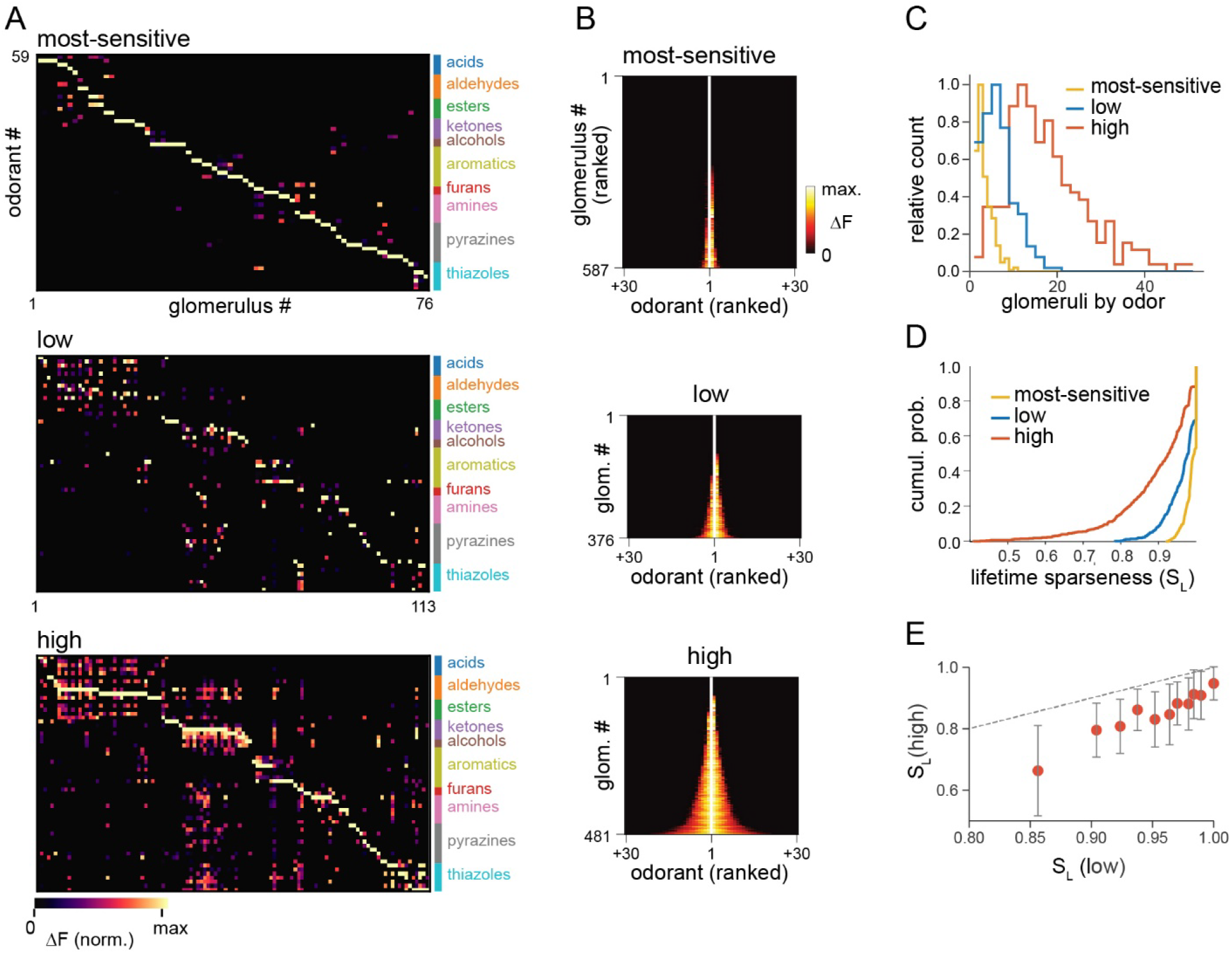
Glomeruli show variable broadening of tuning with increasing odorant concentration. **A.** Response spectra of glomeruli from a single OB to 59 odorants, presented at the ‘most-sensitive’ (top), ‘low’ (middle), and ‘high’ concentration regimes (bottom). Matrices are column-normalized to the maximum response for each glomerulus. Column indexing is sorted to optimize visualization of tuning specificity. Indexing is the same for the ‘low’ and ‘high’ matrices, such that corresponding columns are from the same glomerulus (in the same OB). **B.** Response spectra of all glomeruli pooled across subjects (rows) for each concentration regime, sorted by tuning breadth (lifetime sparseness). Odorants (columns) are sorted by relative response amplitude within each glomerulus (rows). **C.** Probability distributions of the number of glomeruli responding to a given odorant, compiled across the odorant panel and across preparations for the three concentration regimes. **D.** Cumulative probability distributions of lifetime sparseness values for all glomeruli, compiled across the three concentration regimes. **E.** Comparison of lifetime sparseness (S_L_) values within glomeruli evoked by odorants presented at ‘low’ versus ‘high’ concentrations. Measurements were binned based on their ‘low’ concentration S_L_, with all values <1 binned into deciles, and all glomeruli with S_L_ =1 in their own bin. Points indicate mean ± st. dev. of the ‘high’ concentration S_L_ values for the corresponding glomeruli in each bin. Dashed line: unity. S_L_ values at ‘high’ concentrations are lower, as expected, but show substantial variability.

Concentration-dependent changes in tuning markedly varied across glomeruli, however. Many glomeruli remained narrowly tuned even at higher concentrations, with 31% and 12% of all glomeruli responding to only 1 of the 59 odorants (S_L_ = 1) at the ‘low’ and ‘high’ concentrations, respectively (**Fig. 2C,D**). The drop in S_L_ from low to high concentration was highly variable across glomeruli, with a standard deviation of 0.09 (**Fig. 2E**).

Such heterogeneous broadening of glomerular tuning suggests that increasing odorant concentrations may alter the structure of how glomeruli represent odorants across chemical space. In the ‘most-sensitive’ regime, glomerular representations are sparse and high-dimensional but show some structure organized by key chemical features of odorants (Burton et al., 2022). This organization was still apparent after reanalysis of the 59-odorant subset of the ‘most-sensitive’ dataset (**Fig. 3A,B**). Specifically, the correlation matrix of glomerular response vectors (averaged across OBs) exhibited a few blocks of highly-correlated glomerular representations corresponding to thiazoles, amines, and aliphatic acids, esters and aldehydes of the same carbon chain length, embedded in an otherwise sparse matrix consisting of near-zero odorant correlations (**Fig. 3B**). Odorant representations at the ‘low’ and ‘high’ concentrations were less sparse and included larger blocks of higher correlation (**Fig. 3C,D**), collectively reflecting a lower dimensionality of glomerular representational space. Effective dimensionality decreased by nearly half from ‘most-sensitive’ to ‘low’ to ‘high’ concentrations (23.7 vs. 19.3 vs. 13.0).

**Figure 3.**
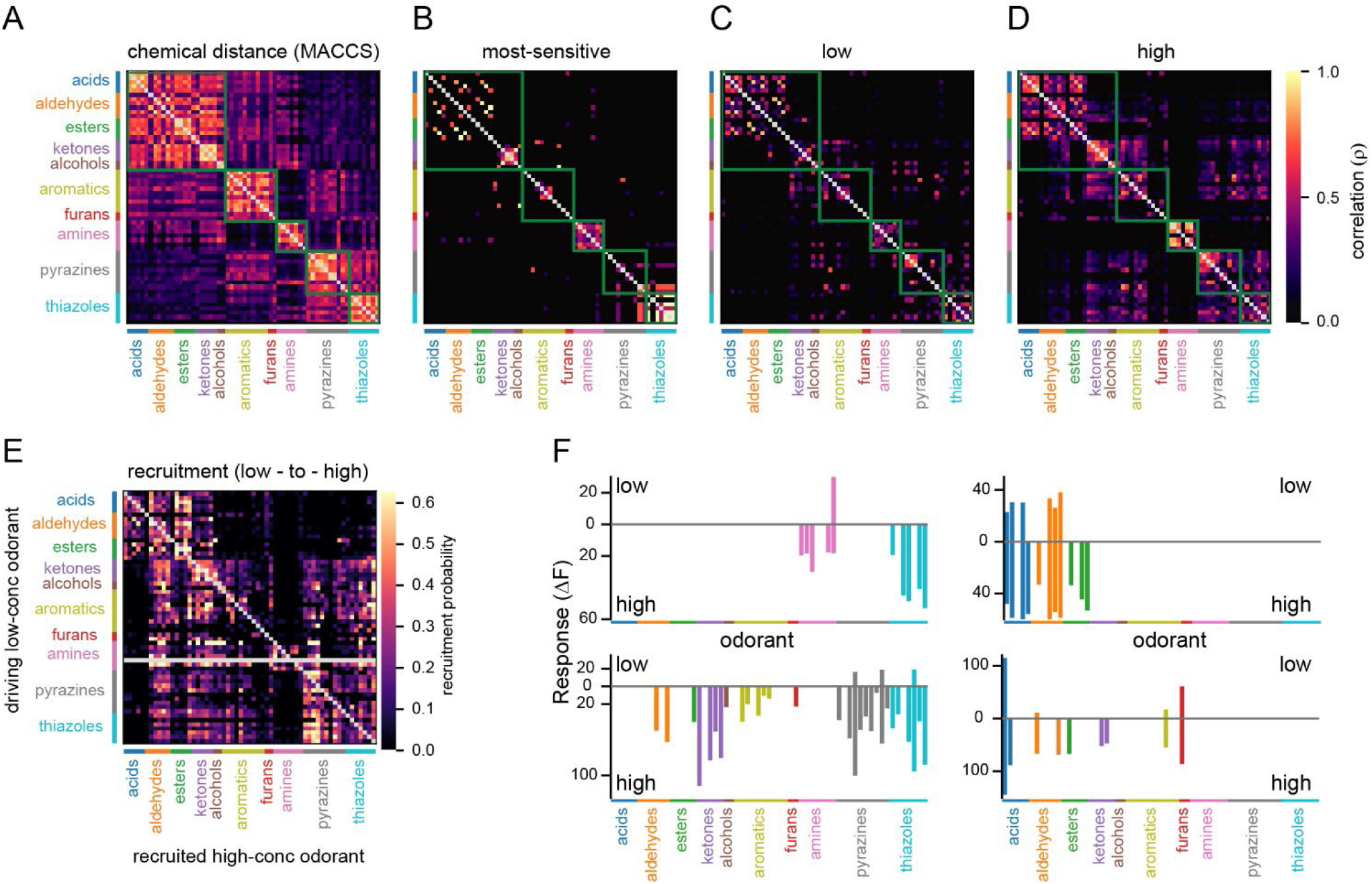
Emergent correlational structure in odor representations reflects selective expansion of response spectra with respect to odorant chemical classes. **A.** Similarity matrix for the 59 odorants, calculated from the cosine distance of the MACCS fingerprints of the odorant panel. Similarity scales from 0 (no similarity) to 1 (identical). Green boxes highlight the most visually salient areas of higher similarity, and correspond to broad chemical classes (from left to right: aliphatic hydrocarbons, aromatic and cyclic hydrocarbons, amines, pyrazines and thiazoles). **B, C, D.** Mean correlation matrices of glomerular response vectors across the odorant panel, computed for each concentration regime. Each value shows the Spearman’s rank correlation (ρ) between the vectors of glomerular responses evoked by a pair of odorants, then averaged across all preparations for each odorant pair. Green boxes are replications from (A) for reference. Note a partial correspondence of the structure in response correlational space to that of chemical feature space, which is more evident at higher concentrations. **E.** Recruitment matrix showing the probability of a glomerulus acquiring responsiveness to an odorant presented at ‘high’ concentration (columns), as a function of its responsiveness to a different odorant at ‘low’ concentration (rows). See Methods for details. Gray row represents undefined values (i.e., no responses at ‘low’ concentration). **F.** Comparison of response spectra for four example glomeruli across the 59-odorant panel at ‘low’ and ‘high’ concentrations. ‘High’ concentration response axis is inverted to mirror the ‘low’ concentration responses for easy visual comparison. Each glomerulus acquires responsiveness to additional odorants at ‘high’ concentration but only among odorants of particular chemical classes.

The emergent structure in the glomerular representational space of higher odorant concentrations appeared to partially map to chemical features of odorants. A distance matrix of chemical similarities among the 59 odorants, based on a binary fingerprint of 166 structural features (MACCS) (Durant et al., 2002) showed a block structure that partially overlapped with the structure of odorant response correlations (**Figs. 3A,D**). For example, responses among the heterocyclic thiazoles and pyrazines became more similar at higher concentrations, recapitulating the structural similarity among these odorants (compare **Figs. 3C,D**). Other emergent patterns of response similarities were selective among subsets of aliphatic or aromatic odorants - for example, carboxylic acids, and aldehydes (but not ketones) - that were not well-distinguished by chemical features (**Fig. 3D**).

We explicitly visualized patterns of concentration-dependent broadening of tuning by generating a ‘recruitment matrix’ that captured the probability of a glomerulus acquiring responsiveness to an odorant at ‘high’ concentration, as a function of its responsiveness to a different odorant at ‘low’ concentration (**Fig. 3E**). Selectivity in concentration-dependent broadening was clearly apparent in the recruitment matrix, with notable features including the expansion of tuning among acid-and ester-sensitive glomeruli to include other acids, esters and aldehydes but no other chemical classes, and the expansion of responsiveness to heterocyclic odorants (e.g., pyrazines and thiazoles) to include a range of other aliphatic hydrocarbons but *not* carboxylic acids. Selective recruitment was also clear in the response spectra of individual glomeruli (**Fig. 3F**), including the somewhat exceptional tuning of glomerulus 155 (see below). Thus, glomerular odorant representations expand *selectively* and *asymmetrically* with respect to chemical feature space as odorant concentrations increase, with responses retaining appreciable chemical class selectivity even at relatively high concentrations.

### Structured tuning to chemical features by the glomerular population persists across odorant concentrations

To better understand the concentration-dependent restructuring of odorant representations, we began by classifying glomeruli according to their tuning at ‘low’ concentrations, then asked how this tuning changed across the same odorant panel presented at ‘high’ concentrations. We used an unbiased, quantitative approach – non-negative matrix factorization (NMF) – to identify clusters of glomeruli with highly correlated odorant response spectra (see Methods). We used NMF rather than principal components analysis because it describes the response of each glomerulus as a sum of non-negative tuning components, such that each component can be decomposed into a group of glomeruli activated by a common set of odorants. The lack of suppressive responses among OSNs further supported the application of NMF to our datasets. We initially excluded glomeruli that responded only at ‘high’ concentration, representing approximately 20% of all glomeruli, as these were uninformative about concentration-dependent changes.

NMF analysis indicated that seven clusters of glomeruli adequately captured the variance in response spectra across the glomerular population; increasing cluster number captured minimal additional variance compared to shuffled data (**Fig. S1AB**), and partitioning beyond seven clusters tended to result in decreasing cluster reproducibility (**Fig. S1C**). The seven clusters were readily apparent in a correlation matrix of glomerular response vectors ordered by cluster membership (**Fig. 4A**). This functional similarity could be directly traced to the responsiveness of glomeruli within each cluster to a characteristic subset of odorants, which were largely organized by chemical class (**Fig. 4B**). While clustering was performed on a concatenated dataset of glomeruli pooled across all four OBs, clusters with comparable tuning were also recovered separately from each OB (**Fig. 4C, Fig. S2**), consistent with their reflecting reproducible features of the OSN repertoire rather than an artifact of pooling across animals. Including the ∼20% of glomeruli that only responded at ‘high’ odorant concentration further yielded 7 clusters with nearly the same membership of glomeruli (with the addition of the high-concentration-recruited glomeruli) (**Fig. 4D**), consistent with the identified clusters robustly capturing distinct tuning profiles of glomeruli that do not depend on the choice of absolute odorant concentration.

**Figure 4.**
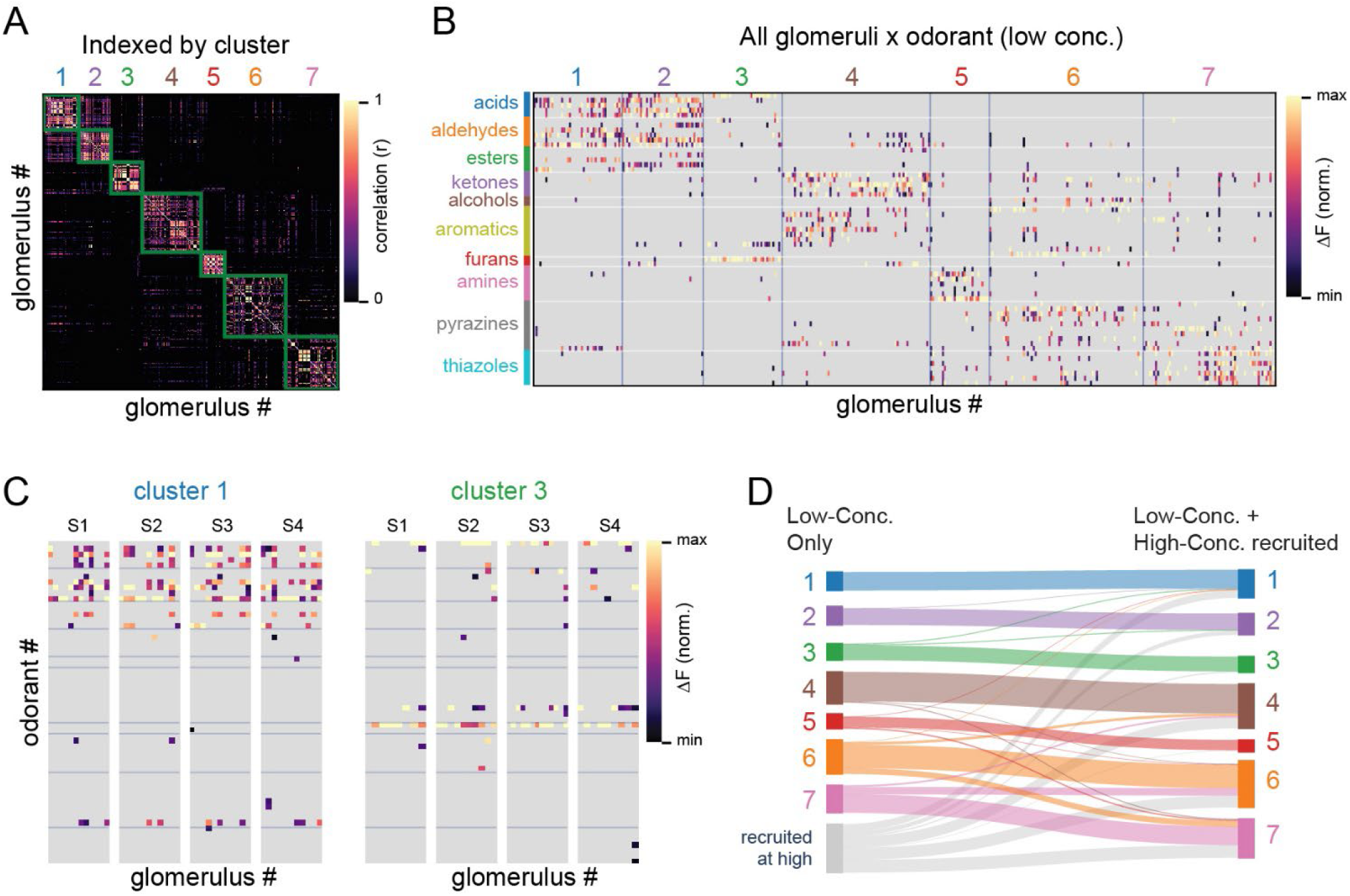
Glomeruli cluster into distinct subpopulations on the basis of shared responses across the ‘low’-concentration odorant panel. **A.** Correlation matrix of glomerulus x glomerulus odorant response vectors (Pearson’s r) for all glomeruli (n = 376, pooled across 4 OBs), with glomeruli sorted by their NMF cluster identity. Green blocks demarcate clustered groups. Within-cluster mean correlation coefficient, 0.12–0.51 (range), between-cluster mean = −0.03. **B.** Glomerulus x odorant response matrix for all glomeruli across the panel of 59 odorants at ‘low’ concentration. Same data and normalization as in Fig. 2A, with glomeruli sorted according to NMF cluster membership. **C.** Glomerulus x odorant response matrices for all glomeruli in two clusters, shown separately for each of the four OBs (subjects “S1-S4”), illustrating consistency of cluster tuning. **D.** Sankey plot of glomerular cluster membership when clustering on ‘low’-concentration responses alone (left) and when including glomeruli responding only to the ‘high’ concentration panel (right). Cluster membership is relatively insensitive to the inclusion of lower-sensitivity glomeruli: most glomeruli retain the same membership, and glomeruli recruited only at ‘high’ concentration distribute to all 7 clusters.

To examine the tuning of glomerular subpopulations with respect to chemical features and concentration dependence, we plotted the fraction of glomeruli in a cluster activated by each of the 59 odorants at ‘low’ concentration and compared this response distribution to that computed for the same glomeruli at ‘high’ concentrations (**Fig 5A**). Glomerular clusters largely mapped to distinct chemical classes of odorants, as further shown in plots of the fraction of glomeruli in a cluster activated by any odorant in each class (**Fig. 5B**). In particular, clusters 1 and 2 were selective for aliphatic acids, aldehydes and, to a lesser degree, esters; cluster 4 was selective for aliphatic and aromatic ketones, cluster 5 was selective for amines, and clusters 6 and 7 were tuned to the heterocyclic pyrazines and thiazoles, respectively (**Fig. 5B**). These results are largely in agreement with those characterizing the tuning of glomeruli at the ‘most-sensitive’ concentration regime, which found that the co-tuning of glomeruli to different odorants is well-predicted by shared structural motifs (Burton et al., 2022).

**Figure 5.**
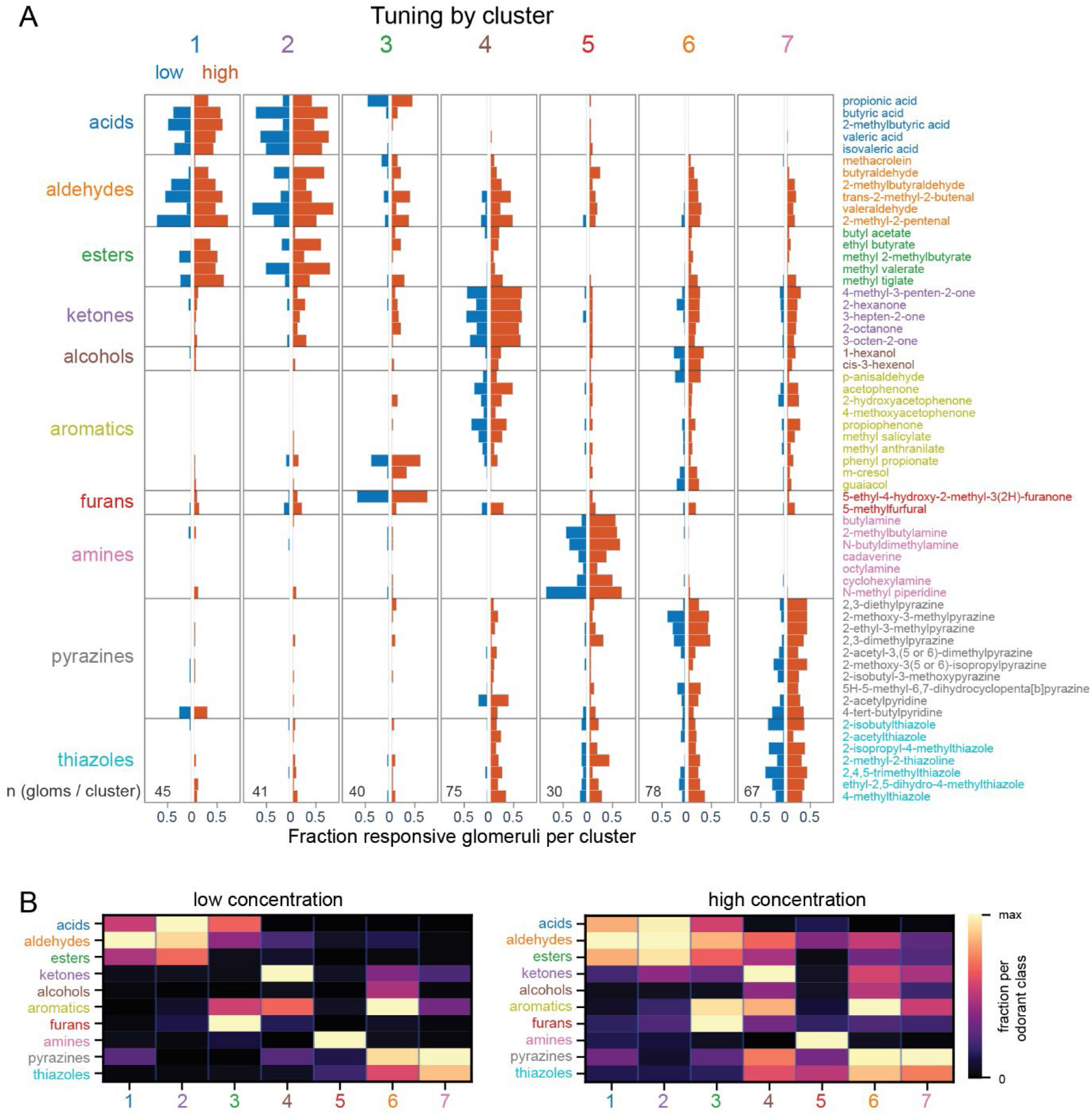
Glomerular clusters respond to odorants within select chemical classes at ‘low’ and ‘high’ concentrations. **A.** Glomerular cluster responsivity across the 59-odorant panel, expressed as the fraction of all glomeruli within a cluster that show a nonzero response to each odorant (response amplitudes are ignored). Blue bars, ‘low’ concentration panel, orange bars, ‘high’ concentration panel. ‘Low’ and ‘high’ concentration response fractions are reflected across the y-axis to facilitate visualization. Odorants are colored to indicate chemical class. Total number of glomeruli in each cluster indicated at bottom of each plot. **B.** Responsivity of glomerular clusters among each chemical class, for ‘low’ and ‘high’-concentration panels. Values plotted are the fraction of all glomeruli within each cluster responding to any odorant in each class, as in (A), column-normalized to the maximum fraction within each cluster.

Glomerular cluster response specificity largely persisted across concentration, such that effective odorants at ‘low’ and ‘high’ concentrations remained the same. At the same time, as seen for individual glomeruli (i.e., Fig. 3), concentration-dependent broadening of cluster responses was not only selective but also asymmetric. For example, glomeruli sensitive to amines at ‘low’ concentration (i.e., cluster 5) show some broadening of their responsiveness to include select heterocylic compounds (thiazoles and pyrazines) at ‘high’ concentration, but glomeruli sensitive to ‘low’ concentrations of heterocyclics (i.e., clusters 6 and 7) do not acquire responsiveness to amines (**Fig. 5A, B**). Likewise, glomeruli in clusters 1 and 2 were distinguished by their responsiveness to carboxylic acids, esters and aldehydes at low concentration; glomeruli in other clusters acquired responsiveness to aldehydes at high concentration but remained selective and unresponsive to the carboxylic acids and, to a lesser degree, esters (**Fig. 5A,B**). These results suggest that the presence of tuning-defined glomerular clusters at low odorant concentrations meaningfully reflects functionally distinct subpopulations of glomeruli that show differential broadening of tuning at higher concentrations.

### Tuning-defined glomerular subsets are spatially organized and align with the projection domains of receptor subfamilies

We found earlier that, in the ‘most-sensitive’ concentration regime, glomeruli with the highest sensitivities to odorants of particular chemical classes are heterogeneously distributed on the dorsal OB surface (Burton et al., 2022). To determine if glomerular subsets defined by their tuning across the 59 odorants presented at higher concentrations were also heterogeneously distributed we mapped the position of glomeruli from each NMF cluster, referenced to the posterior and medial extent of the dorsal OB as in (Burton et al., 2022). This mapping revealed a spatially restricted distribution of glomeruli for each of the clusters (**Fig. 6A, Fig. S3**). Restricted glomerular mapping was particularly evident for the acid-sensitive clusters 1 and 2, with glomeruli almost entirely confined to an anterior-medial wedge of the OB. This spatial organization recapitulates the known projection domain of OSNs expressing class I ORs, which appear specialized to detect carboxylic acids (Bozza et al., 2009; Cichy et al., 2019; Billesbølle et al., 2023). Glomeruli in the amine-sensitive cluster 5 were preferentially located in the central-medial OB, in the region corresponding to the projection domain of TAAR-expressing OSNs (Pacifico et al., 2012). Glomeruli in other clusters, including clusters 4, 6, and 7 (selectively responsive to ketones and/or heterocyclic odorants at ‘low’ concentration), were almost entirely excluded from the apparent class I zone (**Fig. 6A**), and appear to recapitulate the known projection domain of OSNs expressing class II ORs (Bozza et al., 2009).

**Figure 6.**
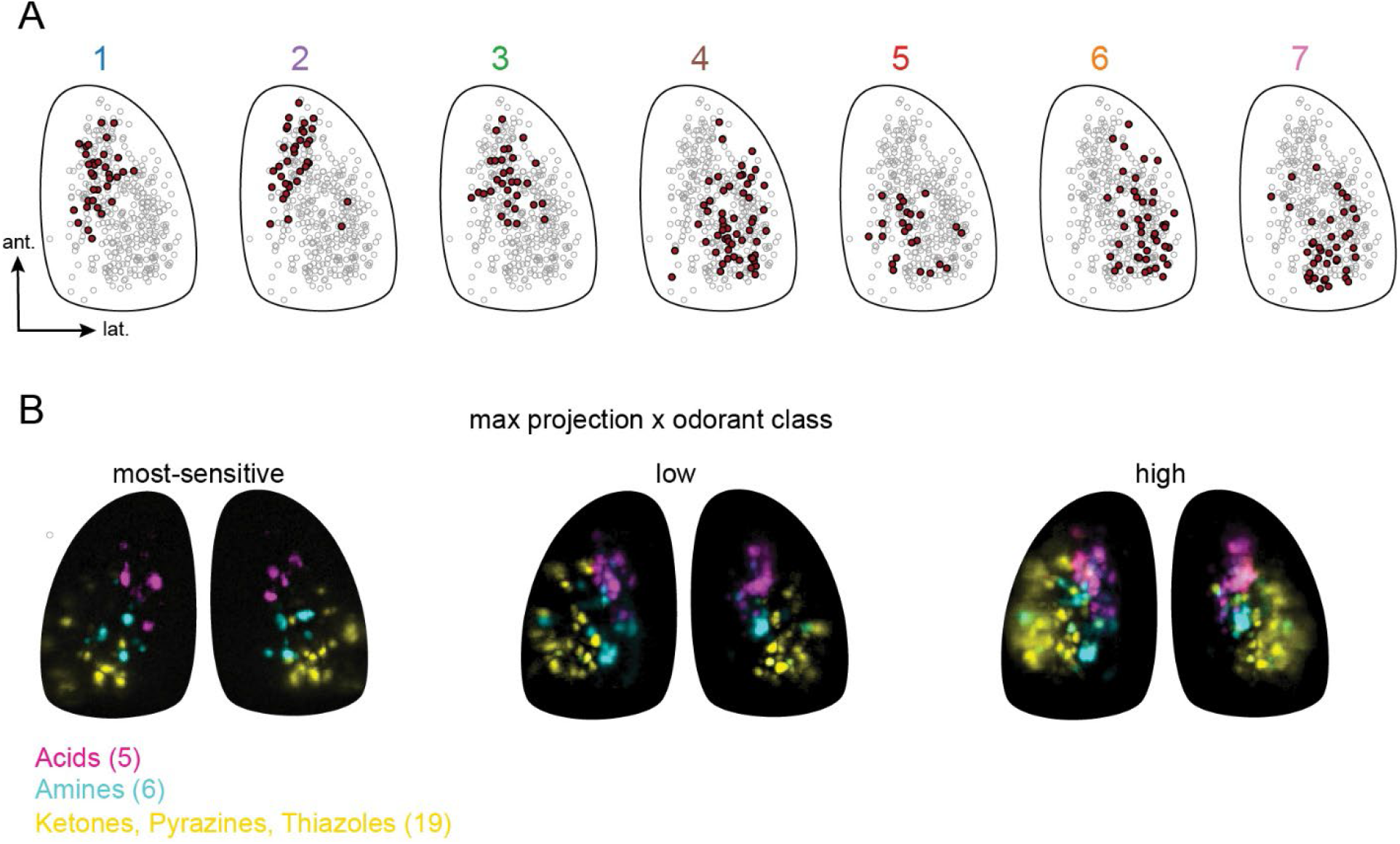
Tuning-defined glomerular clusters align with the projection domains of receptor subfamilies and maintain domain specificity across concentration. **A.** Locations of glomeruli imaged across all 4 OBs (grey) after aligning fields of view (see Methods), with each glomerular location denoted with a circle, and glomeruli of each cluster (number noted above) shown in red. **B.** Maximal projections of ΔF response maps elicited by odorants within distinct chemical classes: carboxylic acids (magenta), amines (cyan), and ketones, pyrazines and thiazoles (yellow), overlaid on the same reference OB outline. ‘Low’ and ‘High’ overlays are from the same mouse. ‘Most-sensitive’ overlay corresponds to mouse 4 from (Burton et al., 2022). Numbers indicate number of odorant response maps overlaid in each group. Increasing concentration recruits more glomeruli but with almost no co-tuning or spatial overlap among groups.

The specificity and spatial segregation of glomeruli with different chemical tuning was especially apparent when mapping responses to distinguishing odorant classes in the same OB. Maximum projections of response maps to all carboxylic acids, all amines, and all ketones and heterocyclics at the ‘most-sensitive’, ‘low’ and ‘high’ concentrations highlighted distinct spatial domains and nearly complete lack of glomerular response overlap (**Fig. 6B**). Higher odorant concentrations activated more glomeruli, as expected, but recruited glomeruli remained within their respective putative receptor-defined domains. This spatial examination, together with our earlier analysis of glomerular cluster responsivity (**Fig. 5**), strongly suggests that the three major receptor subfamilies – class I ORs, class II ORs, and TAARs – are primary organizing features of the tuning structure of odorant representations across the OSN repertoire as well as the topography of response patterns in the OB. Notably, this organization persists across at least 3-4 log units of concentration.

### Concentration-dependent broadening of glomerular tuning is predictable, selective and structured according to consensus tuning of OSN subpopulations

Thus far our analysis reveals that, rather than a uniform broadening with increasing odorant concentrations, glomerular tuning broadens in a manner that is selective and asymmetric across chemical feature space and appears organized along functionally-distinct subpopulations of glomeruli. To more rigorously explore these qualitative observations, we developed a statistical approach to characterize how the tuning of glomeruli at ‘low’ concentration predicts their change in tuning across odorants at ‘high’ concentration. We used partial least squares regression (PLS-R) (Wold et al., 2001; Krishnan et al., 2011) to calculate a common weighting matrix (*W*) that relates the ‘low’-concentration glomerular response vector for each odorant (*X*) (i.e., a 1 odorant x 376 glomerulus vector, concatenated across OBs) to the ‘high’-concentration vector for the same odorant (*Y*), or *Y = XW* (**Fig. 7A**). Compared to a conventional regression strategy, PLS-R reduces the risk of overfitting by identifying a few latent components (i.e., patterns of shared glomerular tuning) that account for a large portion of the covariance between ‘low’-and ‘high’-concentration responses, in contrast to direct regression, which would involve predictions for a smaller number of samples (i.e., the 59 odorants) from a larger population of predictors (i.e., the 376 glomeruli, yielding 376 × 376 ≈ 141,000 pairwise relationships). In our implementation (see Methods), we fixed the PLS-R model at eight latent components.

**Figure 7.**
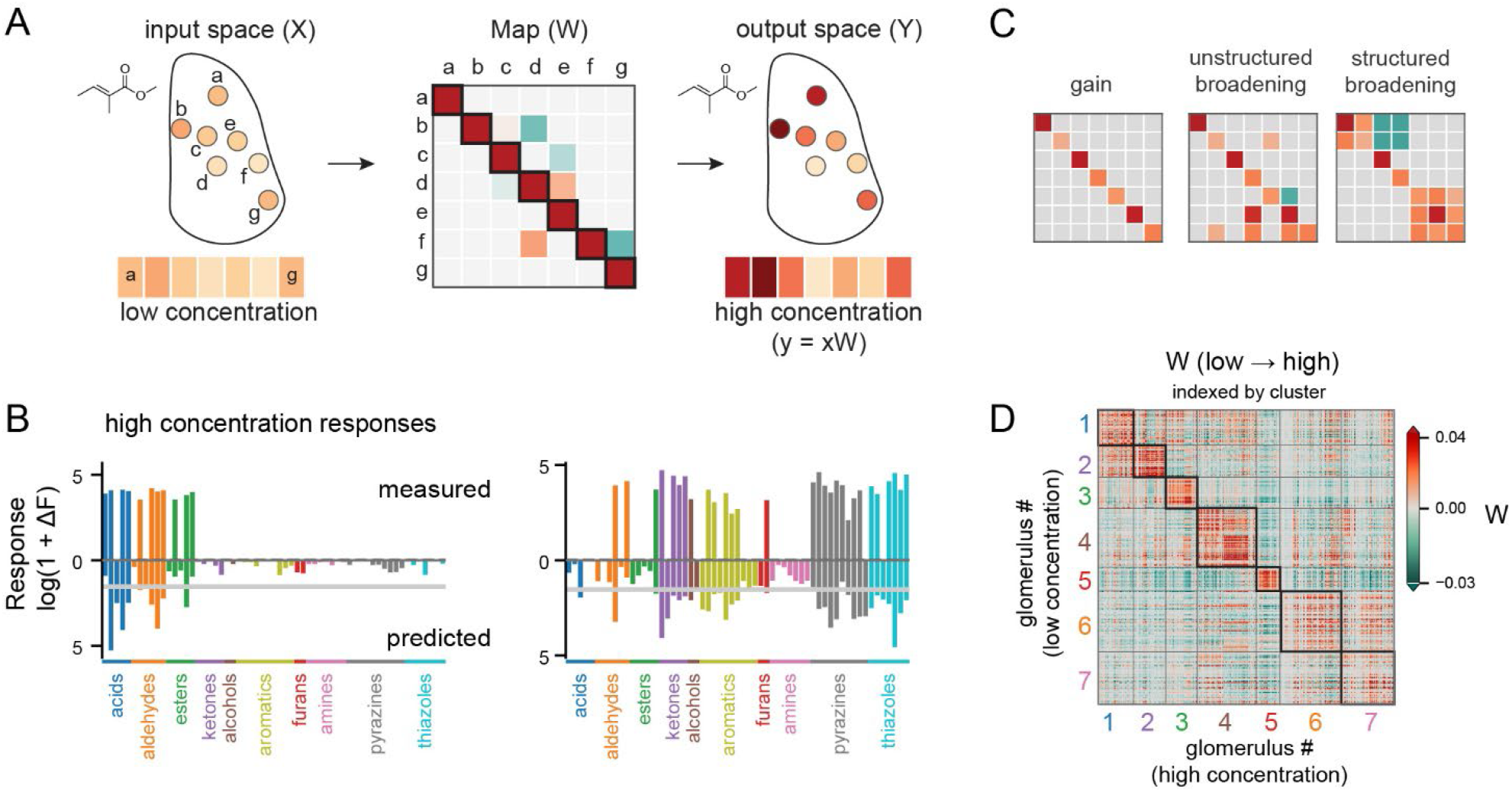
Predicting high-concentration response spectra from low-concentration responses using partial least-squares regression reveals structured concentration-dependent broadening rules. **A.** Schematic of PLS-R strategy for predicting glomerular tuning at ‘high’ concentrations from ‘low’-concentration responses. **B.** Plots of measured (top half) and predicted (lower half) high-concentration response spectra for two example glomeruli (same as in Fig. 3), plotted on a log scale. All predictions are out-of-sample. Gray horizontal line indicates lowest observed ΔF value attributed to an odorant response across the experimental datasets. **C.** Schematics representing three possible PLS-R weight matrix configurations depending on the nature of concentration-dependent recruitment of glomerular responses. See Text for details. **D.** Full PLS-R glomerulus x glomerulus weight matrix (fitted once to all 59 odorants) describing ‘low’-to ‘high’-concentration changes in responses, with glomeruli indexed by their NMF cluster memberships.

To evaluate the effectiveness of PLS-R in predicting ‘high’-from ‘low’-concentration responses, we calculated the weighting matrix *W* on a subset of data and tested prediction accuracy on held-out odorants under 4-fold cross-validation, stratified by chemical class to ensure that each evaluation of *W* reflected the chemical diversity of the panel. Using this approach, the model explained 27% of the variance in high-concentration response amplitudes. While this performance may seem modest, we note that the model was evaluated on its prediction of response amplitudes (as opposed to simply binary responsiveness), based on ‘low’-concentration responses that were themselves extremely sparse.

Examples of held-out predicted and recorded responses to the ‘high’ concentration odorant panel, taken from the same examples shown in **Fig. 3F**, are shown in **Fig. 7B**. While the model predicted more broadly-tuned responses than were seen in the experimental data, incorrectly-predicted non-zero responses tended to be smaller in amplitude than the minimum measured experimental response (**Fig. 3F**, **Fig. S4**) (note that the example comparisons are plotted on log scale).

The matrix *W*, calculated through PLS-R across all odorants, effectively contains a recipe for both scaling and combining ‘low’-concentration glomerular response spectra to predict changes in their response spectra at ‘high’-concentration. Elements along the diagonal of *W* strictly describe within-glomerulus scaling (i.e., a ‘gain’ function) while off-diagonal elements of *W* describe how tuning broadens at higher concentrations. To gain insight into the logic of this broadening, we next examined the structure of these off-diagonal elements.

If glomerular tuning broadens randomly with respect to odor space and with respect to other glomeruli, off-diagonal weights should be unstructured (**Fig. 7C**). Instead, off-diagonal weights emerged in structured blocks with respect to glomerular identity. In particular, ordering the rows and columns of *W* according to the membership of each glomerulus in the functional glomerular clusters defined earlier by NMF revealed a clear block organization in the matrix *W* that aligned with the blocks of cluster membership (**Fig. 7D**). Quantitatively, the block-diagonal region comprising within-cluster weights accounted for 76% of the predictive power of the matrix *W*. This result supports the conclusion that concentration-dependent broadening is heterogenous across the OSN population, with glomeruli exhibiting similar ‘low’-concentration tuning (i.e., glomeruli within an NMF cluster) undergoing more similar broadening of their receptive fields than glomeruli exhibiting different ‘low’-concentration tuning (i.e., glomeruli across distinct NMF clusters).

We further tested this conclusion by asking whether glomeruli with a given ‘low’-concentration consensus tuning profile (i.e., NMF cluster) broaden their tuning at ‘high’ concentrations by incorporating consensus tuning features of glomeruli from *select* other clusters. To do so, we used the mean ‘low’-concentration tuning profiles of the seven NMF-defined clusters to define a set of canonical tuning motifs – a compact library from which any given glomerular tuning profile could be approximated through linear combination (**Fig. 8A**). For each glomerulus, we first asked how well its ‘high’-concentration response could be predicted by a simple gain-like scaling of the canonical motif associated with its own cluster (i.e., ‘source’). We then tested, one at a time, whether adding the tuning motif of each of the other six clusters improved prediction of ‘high’-concentration responses for held-out odorants (measured as the improvement in cross-validated R^2^; see methods). This allowed us to distinguish simple broadening within a glomerulus’s own consensus tuning motif from selective broadening of tuning toward odorants more characteristic of other cluster motifs.

**Figure 8.**
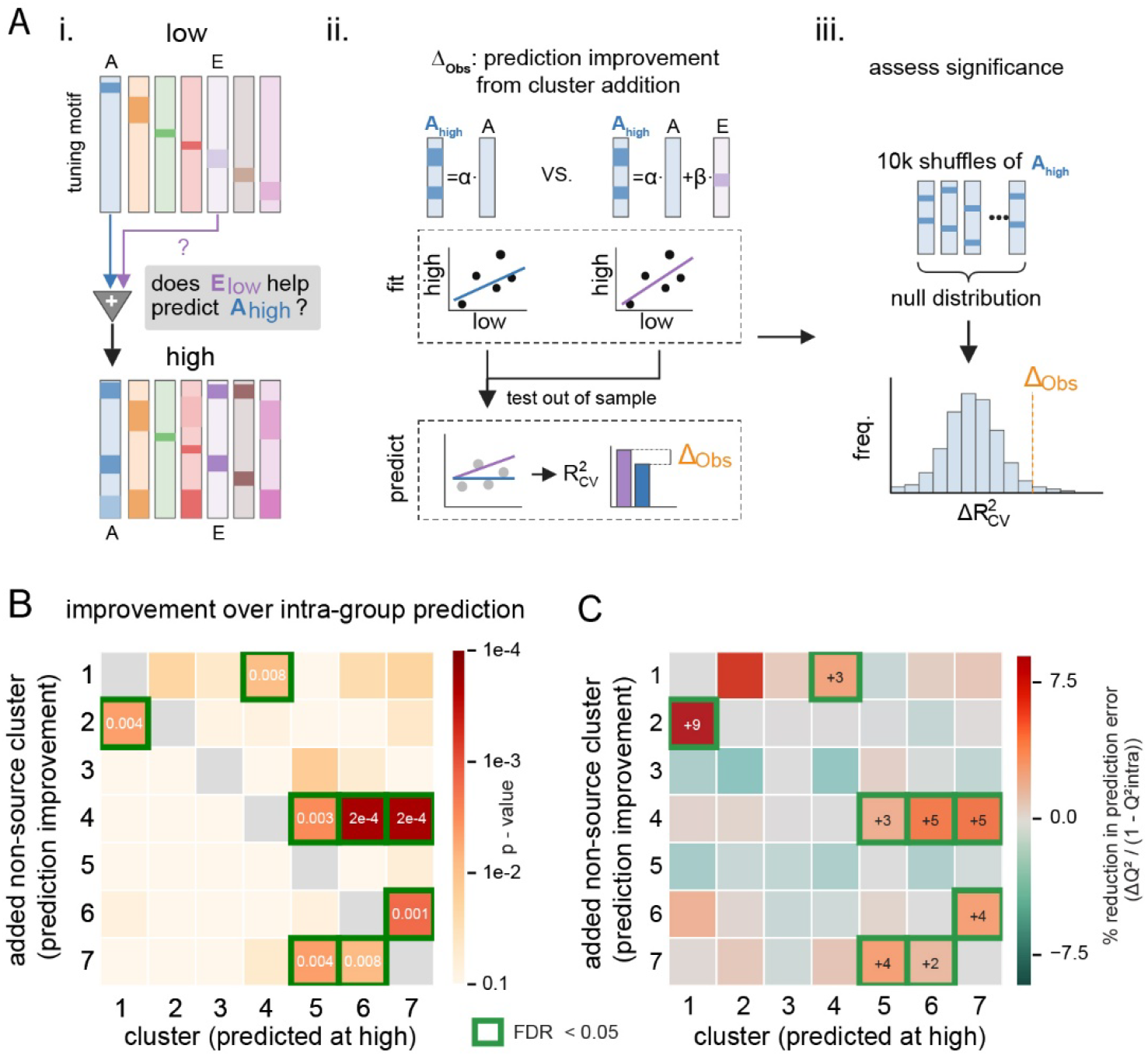
Concentration-dependent broadening can be described by selective acquisition of responsiveness to glomerular tuning motifs. **A.** Schematic of test for cluster-based prediction of concentration-dependent broadening. Each glomerulus’s high-concentration responses are predicted from the mean low-concentration response profile (‘tuning motif’) of its own NMF cluster, and also from that motif plus the motif of one other cluster; the improvement in predicting held-out odorants is compared with a null distribution (see Methods). **B.** Statistical significance of the prediction improvement achieved by adding the tuning motif of a cluster (rows, ‘non-source cluster’) to that of each source cluster (columns). Heatmap shows p-values against a null distribution of 10,000 random cluster pairings. Outlined pairs are significant after Benjamini–Hochberg correction (FDR < 0.05). **C.** Effect size for the same pairs in (B): the percent reduction in the source cluster’s prediction error when the non-source cluster’s motif is added.

We aggregated improvements in prediction for each non-source cluster, yielding 42 possible directional relationships between a source and non-source motif. Across all 42 directional pairs, eight relationships proved significant at a false discovery rate (FDR) < 0.05 (against a null distribution of 10,000 permutations randomly pairing ‘high’ and ‘low’-concentration responses, followed by Benjamini–Hochberg - correction, see Methods) (**Fig. 8B**). **Fig. 8C** shows the effect size of each relationship (significant pairs outlined), expressed as the percent reduction in prediction error when the non-source tuning motif was added to the source tuning motif to predict ‘high’-concentration responses. Notably, the presence of select and asymmetric relationships revealed in our earlier analyses of odorant recruitment (**Fig. 3E**) and fractional cluster activation (**Fig. 5**) was substantially conserved. For example, glomeruli sensitive to amines at ‘low’ concentration (i.e., **Fig. 8B** matrix column 5) broadened their tuning by acquiring responsivity similar to that of clusters 4 and 7 (i.e., matrix rows 4 and 7), encompassing responses to ketones and heterocyclics, while no clusters broadened their tuning by acquiring responsivity similar to that of cluster 5 (i.e., responses to amines). This analysis suggests that a few canonical response motifs across chemical space can capture heterogeneity in the tuning of glomerular subpopulations at a given concentration, as well as the selective broadening of this tuning at higher concentrations.

## DISCUSSION

Understanding how the nervous system represents the high-dimensional space of olfactory stimuli is a longstanding problem that has been difficult to address due to the nature of olfactory stimulus space and a lack of consensus on the tuning properties of olfactory receptors and their cognate OSNs. OSNs have been variously described as narrowly or broadly tuned and their tuning varies with odorant concentration (Sato et al., 1994; Duchamp-Viret et al., 2000; Hallem and Carlson, 2006; Pelz et al., 2006; Saito et al., 2009; Nara et al., 2011; Ma et al., 2012; Burton et al., 2022). The present results expand on previous studies and are unique in that they characterize the tuning of a relatively large subset (∼10%) of the OSN repertoire using a diverse and reasonably large panel of odorants across concentrations ranging from near-threshold for activating the most-sensitive OSNs to 3 - 5 log units more concentrated. This comparison revealed several key features of OSN tuning, concentration dependence, and the broader representation of olfactory stimulus space.

First, we find that OSN tuning broadens selectively and heterogeneously across concentration, with OSNs retaining selectivity to particular chemical features even as they acquire responsiveness to new odorants. Second, we find that this concentration-dependent broadening is not only selective across chemical feature space but predictable for OSN subpopulations that are functionally defined on the basis of their sensitivity to lower-concentration odorants. Third, we find that a lower-dimensional structure in the representation of olfactory chemical space emerges with increasing concentration. These conclusions emerged from three complementary analyses of our data: observation of odorant-specific patterns of recruitment that were consistent across glomeruli (i.e., **Fig. 3**); the grouping of glomeruli into clusters defined by distinctly selective tuning across the 59-odorant panel at ‘low’ concentration, (**Figs. 4 and 5**), and a regression model that predicted ‘high’ - concentration response spectra from the ‘low’-concentration responses at the level of both individual glomeruli and glomerular clusters (Figs. 7 and 8). Notably, this lower-dimensional structure further reflected major olfactory receptor subfamilies and their tuning to distinct chemical classes of odorants across a large range of concentrations, apparent in the close spatial alignment of tuning-defined glomerular clusters with putative receptor domains (**Fig. 6**). We discuss implications of these findings below.

### Selective concentration-dependent broadening of OSN tuning

A common view of olfactory receptors and their cognate OSNs is that they are specific but broadly tuned, showing high sensitivity to a few odorants but losing their selectivity at higher concentrations (Sato et al., 1994; Duchamp-Viret et al., 2000; Wachowiak and Cohen, 2001; Hu et al., 2020). Our results suggest a more nuanced view of receptor tuning, in which responsiveness broadens to include additional odorants while maintaining selectivity for specific chemical features. While glomeruli (representing OSN populations expressing a given receptor) showed broadened tuning with increasing odorant concentration, lifetime sparseness – a measure of response selectivity – remained remarkably high overall, suggesting that OSNs remain narrowly tuned even as odorant concentrations increase. This conclusion is consistent with a few prior studies that have probed OSN tuning across a broad array of odorants (Nara et al., 2011), and with an emerging understanding of the structural basis for selectivity of receptors to chemical features (del Mármol et al., 2021; Billesbølle et al., 2023; Guo et al., 2023).

### Mapping glomerular subpopulations to olfactory receptor subfamilies

Our results suggest that much of the structure and heterogeneity of OSN odor representations reflects the major subfamilies of olfactory receptors: class I ORs, class II ORs, and TAARs. Functionally-defined glomerular clusters mapped to spatially distinct regions of the dorsal OB that aligned with the projection domains of the three receptor subfamilies (Bozza et al., 2009; Pacifico et al., 2012; Wang et al., 2022), and their consensus response spectra aligned with previously-established sensitivities of each subfamily to certain odorants (Bozza and Kauer, 1998; Pacifico et al., 2012; Cichy et al., 2019).

The ability to capture glomerular representations of OSN subpopulations that appear to map to defined receptor subfamilies provides validation for the functionally-defined clustering. Notably, other functionally-defined glomerular clusters were spatially restricted within the class II OR-projection domain – for example, clusters 6 and 7, with selective tuning for the heterocyclic pyrazines and thiazoles, as reported in some prior studies (Soucy et al., 2009; Matsumoto et al., 2010). Pyrazine-and thiazole-sensitive receptors, identified in vitro (Marcinek et al., 2021), have OSN projections to regions of the dorsal OB aligned with the locations of cluster 6 and 7 glomeruli (Bintu et al., 2026). It will thus be interesting to determine whether these or additional functionally-defined glomerular clusters map to receptor subfamilies within the class I or class II OR domains. Coupling recent advances in mapping receptors to their cognate glomeruli (Wang et al., 2022; Zhu et al., 2022; Bintu et al., 2026) with the functional characterization of odorant response spectra will be important in addressing this question.

While the sensitivity of class I ORs, class II ORs, and TAARs to ccarboxylic acids, other diverse odorants, and amines, respectively, is well-established, their specificity across concentration – and with respect to each other – has not previously been investigated. The present results suggest that OSNs expressing these different receptor subfamilies remain functionally distinct even at high concentrations. In particular, presumptive class I glomeruli responded to carboxylic acids and their ester and aldehyde derivatives, but remained insensitive to nearly all other compounds even at the highest concentrations tested. The converse was also striking, in that all other glomeruli remained *insensitive* to carboxylic acids even at the highest concentrations tested. Likewise, glomeruli sensitive to amines at low concentration – likely reflecting TAAR-expressing OSNs (Pacifico et al., 2012) – acquired responsiveness to other compounds at higher odorant concentrations, but other (presumptive class I and class II OR-expressing) glomeruli almost never acquired responsiveness to amines. These observations highlight that receptors maintain their specificity as odorant concentrations increase. Recent advances in determining the structure of mammalian olfactory receptors – including TAAR and class I ORs – have focused on understanding the structural basis of ligand recognition (del Mármol et al., 2021; Billesbølle et al., 2023; Guo et al., 2023). Our results suggest that it is equally important to understand the basis for *exclusion* of certain ligands from receptor activation, even when those compounds are present at high concentrations.

### Interpreting odor coding space in different concentration regimes

Given the changes in the structure/dimensionality of odor representation across concentration, it is important to consider how the ‘most-sensitive’, ‘low’ and ‘high’ concentration regimes relate to the occurrence of odorants in the natural environment. Based on a recent meta-analysis of odorant concentrations in various settings (Wachowiak et al., 2025), odorants are most often encountered at concentrations aligned with the ‘most-sensitive’ dataset, where ambient concentrations in natural settings such as a forest and in the volatile headspace of natural odor sources (e.g., fruits, flowers, herbs) are at a range of <1 to ∼ 10 ppb. Odorants in the ‘low’ concentration regime ranged from ∼10 ppb to ∼1 ppm, roughly corresponding to the high end (i.e., 90th percentile) of headspace odorant concentrations or to peak odorant concentrations in high-intensity odor environments. The ‘high’ concentration regime, ranging from ∼1 to >10 ppm, is likely rarely encountered in natural settings, save for highly volatile compounds or possibly while sampling directly from the surface of an odor source. At the same time, however, the ‘high’ regime is aligned with the majority of previous experimental studies characterizing odorant representations in animal models (Wachowiak et al., 2025). Thus, changes in tuning from the ‘most-sensitive’ to the ‘low’ concentration dataset may be most important for understanding how odor representations change from ambient to high-intensity natural settings, while the change in tuning from ‘low’ to ‘high’ odorant concentrations may be more relevant for interpreting earlier studies.

In this regard, earlier studies characterizing odor representations at the level of OB glomeruli generally agree with the results reported here for the ‘high’ concentration panel, finding spatial clustering of glomeruli defined by their responsiveness to particular odorant classes (Uchida et al., 2000; Takahashi et al., 2004; Johnson and Leon, 2007; Matsumoto et al., 2010). Other glomerular mapping studies have observed less specificity in odorant-evoked response patterns, concluding that individual glomeruli or their innervating neurons are broadly tuned, activated by structurally diverse odorants and show little spatial relationship to chemical feature selectivity (Soucy et al., 2009; Ma et al., 2012; Chae et al., 2019). However even these studies observed distinct tuning and spatial localization of glomeruli with sensitivities to acids and aldehydes or to the heterocyclic thiazoles (Soucy et al., 2009; Matsumoto et al., 2010; Ma et al., 2012), located in the presumptive projection domains of class I and II OR-expressing OSNs, respectively, underscoring the prominence of olfactory receptor subfamilies as an organizing feature of odor representations across concentration.

The maintenance of distinct tuning by OSN subpopulations across concentration regimes is consistent with their serving distinct functional roles in olfactory processing. Among the major OR classes, the preferred ligands for class I ORs and TAARs - carboxylic acids and amines - tend to arise from sources with distinct ethological relevance from the more diverse ligands of class II ORs: both are common products of bacterial action and organic decomposition (i.e., spoilage or fermentation) (St. Angelo et al., 1996; Pircher et al., 2007; Ng et al., 2023), and amines can serve as intraspecies signals of sexual maturity or as kairomones signaling predator identity (Dewan et al., 2013; Liberles, 2015; Cichy et al., 2026). In addition, acids and amines have been shown to elicit innate behavioral responses mediated by class I ORs or TAARs (Kobayakawa et al., 2007; Dewan et al., 2013; Liberles, 2015; Cichy et al., 2019; Cichy et al., 2026). Other OSN subpopulations may also be preferentially tuned to odorant classes with distinct meaning and possibly driving distinct behavioral responses - for example, glomeruli tuned to the heterocyclic pyrazines and thiazoles, or terpenoids, which were untested in the current study but which serve as chemical signals arising from plant sources (Gershenzon and Dudareva, 2007).

It is also interesting to consider whether the functional heterogeneity apparent at the level of OSN inputs persists at the level of OB output. Defining input-output transformations in vivo has been difficult due to the high-dimensional nature of the input space (Chae et al., 2019). The robust lower-dimensional structure described in the present study provides a useful framework for better describing these transformations, and for investigating the functional organization of the OB circuits that mediate them. A variety of interneuronal circuits transform sensory input both within and across glomerular channels (Wachowiak and Shipley, 2006; Nagayama et al., 2014; Burton, 2017). Recent evidence suggests that OB circuits mediate selective inhibition among glomeruli (Fantana et al., 2008; Economo et al., 2016), but the determinants of this selectivity are unclear. Interglomerular circuits may be structured to reflect the tuning of glomerular subpopulations, which can serve to decorrelate of overlapping input patterns (Banerjee et al., 2015; Zavitz et al., 2020). Interglomerular inhibition also appears to have a distance-dependent constraint aligned with the spatial distribution of similarly-tuned glomeruli observed here (Luo and Katz, 2001; Lehmann et al., 2016; Dalal and Haddad, 2024). Thus, interglomerular inhibition may preferentially act between mitral/tufted cells linked to similarly-tuned glomeruli that are progressively recruited by increasing odorant concentrations.

### Limitations of the study

Several limitations are important to consider in interpreting the results reported here. Foremost, our analysis was confined to the dorsal OB, reflecting technical constraints of our functional imaging approach. This limitation is shared with many prior studies that have, collectively, contributed to our current understanding of glomerular odor representations in the OB (Belluscio and Katz, 2001; Meister and Bonhoeffer, 2001; Wachowiak and Cohen, 2001; Spors et al., 2006; Vincis et al., 2012), input-output transformations (Storace and Cohen, 2017; Chae et al., 2019), the organization and function of OB circuitry in vivo (Luo and Katz, 2001; Fantana et al., 2008; Banerjee et al., 2015; Economo et al., 2016), and the nature of centrifugal feedback to the OB (Rothermel and Wachowiak, 2014; Boyd et al., 2015; Chae et al., 2022). A central assumption of all of these studies is that general principles uncovered from imaging the dorsal OB generalize across its full extent. For the current study, it is likely that we failed to uncover some aspects of the representational structure of olfactory space, given evidence that certain odorant classes preferentially activate glomeruli in the ventral or lateral OB (Igarashi and Mori, 2005; Johnson and Leon, 2007). However, the dorsal OB represents a useful microcosm of olfactory coding within which OSNs expressing all major olfactory receptor subfamilies converge (Pacifico et al., 2012). We therefore expect our main conclusions to apply across the OB, particularly as they relate to the organization of odorant tuning and concentration dependence by expression of distinct olfactory receptor subfamilies. Similarly, while comparatively large and diverse, our 59-odorant panel nevertheless still covers only a fraction of olfactory stimulus space (**Fig. 1A**). Extending experiments to a larger odorant panel (e.g., our original 185 odorant panel; (Burton et al., 2022)) would be useful to better delineate the heterogeneous structure of odor representations.

## METHODS

### Animals

For the ‘low’ and ‘high’ concentration datasets presented here, data were collected from 2 male and 1 female compound heterozygous crosses of OMP-IRES-Cre (Jackson Laboratory stock #006668) and Rosa26-CAG-lox-STOP-lox-GCaMP6f (RCL-GaMP6f; Ai95D; Jackson Laboratory stock #024105) mouse lines, aged 16 -18 weeks. Mice were housed up to 5 per cage on a 12-hour light/dark cycle with food and water available ad libitum. All procedures were performed following the National Institutes of Health Guide for the Care and Use of Laboratory Animals and were approved by the University of Utah Institutional Animal Care and Use Committee. The ‘most-sensitive’ dataset, originally presented in (Burton et al., 2022), was collected in OMP-IRES-tTA x tetO-GCaMP6s reporter crosses.

### Olfactometry

Odorants were obtained from Sigma-Aldrich, TCI America, Bedoukian Research, or ICN Biomedicals and delivered using a custom olfactometer that allowed mixing of odorant vapor in free space prior and computer-controlled timing and choice of odorant identity (Burton et al., 2019). Odorant delivery parameters were identical to those used in (Burton et al., 2022) except that the end-stage eductor was omitted. Non-amine odorants were diluted in caprylic/capric medium chain triglyceride oil (C3465, Spectrum Chemical Mfg. Corp.) within ∼1 week of experiments, as described in (Burton et al., 2022); amine odorants were freshly diluted in water immediately prior to each experiment to minimize odorant oxidation. Dilutions in the liquid phase ranged from 1:10 - 1:1250 in order to achieve the estimated delivered concentrations reported in the Results.

### Data collection and processing

Odorant responses were collected with epifluorescence imaging in isoflurane-anesthetized mice outfitted with a double tracheotomy to decouple respiration and anesthetic delivery from odorant presentation and sampling, using procedures identical to those described in (Burton et al., 2022). Data was obtained from three mice, and four OBs total. Due to technical issues, responses to the amines were collected in only one of the three mice, from both the left and right OBs. Odorant response matrices (odorant x glomerulus) were generated using the analysis pipeline described in (Burton et al., 2022). Briefly, averaged ΔF response maps were generated from 4 - 6 repeated presentations of each odorant, after which regions of interest (ROIs) were selected manually using custom visualization software written in MATLAB (https://github.com/WachowiakLab). ROIs representing presumed individual glomeruli from the dorsal OB were segmented manually, with the segmenter blind to odorant identity (Burton et al., 2022). To eliminate misattribution of responses due to lateral spread of fluorescence signals from adjacent glomeruli that is inherent in the widefield imaging in scattering tissue, the segmentation procedure involved sorting odorant response maps in descending order of their ΔF response for a given ROI, and setting to zero any entries in the response matrix that did not correspond to clear ΔF signal foci that were centered within the ROI of interest. After segmentation, response matrices were generated using the mean ΔF value across all pixels in an ROI, for all 59 tested odorants.

### Data Analysis

#### Chemical feature space

To visualize odorant relationships we used the Python package RDKit. To visualize and quantify odorant similarities based on physicochemical features (e.g., **Figs. 1A,3A**), a physicochemical feature space was constructed from 4789 odorous compounds in the combined GoodScents/Leffingwell dataset (Lee et al., 2023), as provided by the OpenPOM github site (https://github.com/ARY2260/openpom). This list was supplemented with 31 additional compounds from our experimental odorant panel. The Python package RDKit (https://www.rdkit.org/) was used to generate molecular fingerprints for each compound from its SMILES string; fingerprints were encoded as MACCS keys, a binary representation indicating the presence or absence of 166 chemical features (Durant et al., 2002). Principal component analysis (PCA) embeddings were computed from MACCS fingerprints of the chemical space dataset. For display (**Fig. 1A**), a three-dimensional scatter projection of the first three PCs was visualized using OriginLab (Origin(Pro) 2026, OriginLab Corporation, Northampton, MA, USA).

#### Characterization of glomerular response spectra

Lifetime sparseness (S_L_) and population sparseness (S_P_) measures were computed from ΔF response vectors using a standard formula, as described in (Burton et al., 2022). For both measures, a value of 1 indicates maximally selective tuning (a glomerular response to only one of the 59 odorants (S_L_), or a single glomerulus activated by an odorant (S_P_), while a value of 0 indicates completely nonselective tuning, i.e., equivalent responses to all odorants or equivalent activation of all glomeruli. Response correlation matrices were computed using rank correlation (Spearman’s ρ) across all glomeruli in a response matrix for all odorant pairs; the resulting odorant correlation matrices were averaged across the 4 imaged OBs (8 OBs for the ‘most-sensitive’ dataset). Odorant x odorant recruitment probabilities (**Fig. 3E**) were computed to reflect the probability of a glomerulus acquiring responsiveness to an odorant presented at ‘high’ concentration as a function of its responsiveness to a reference odorant at ‘low’ concentration. Responsiveness (for both reference and recruited glomeruli) was defined as any response greater than 20% of the maximum response of that glomerulus across all odorants. Rows for odorants with no responsive glomeruli have zero values.

#### Functional clustering of glomeruli

Glomeruli were partitioned into tuning clusters by non-negative matrix factorization (NMF; scikit-learn) of the odorant × glomerulus response matrix; a small positive offset (single-precision machine epsilon) was added to inputs to eliminate zero values. We used 7 components (termed ‘canonical’ cluster number; see below), a fixed random seed and 1000 maximum iterations, with default scikit-learn initialization and solver otherwise. Each glomerulus was assigned to the component on which its NMF loading was maximal (hard, argmax assignment). The canonical clustering used for subsequent analyses was computed from the low-concentration responses of the 376-glomerulus set only (yielding 7 clusters), after which the high-concentration matrix was reordered by these low-concentration labels to preserve the low→high correspondence.

The number of clusters was chosen by two criteria, as illustrated in **Fig. S1**. First, the fraction of variance explained by rank-*k* reconstructions was computed for PCA, NMF, and PCA on randomly permuted data (**Fig. S1A**); the permuted-PCA curve, which rises only linearly, provided a noise floor, and the marginal variance explained per added component (the discrete slope; **Fig. S1B**) was compared to this floor to bound the number of components capturing genuine structure. Second, clustering stability was assessed by consensus clustering over 50 random NMF initializations, summarized by the consensus (co-clustering) matrix and by the adjusted Rand index (ARI) of each initialization’s partition to the consensus partition (**Fig. S1C**). Robustness across animals was assessed by computing NMF clusters for each subject independently (**Fig. S2**).

### Partial least-squares regression model

Each odorant’s high-concentration population vector was modeled as a linear transform of its low-concentration vector, Y = XW, with X and Y the paired 59 odorant × 376 glomerulus matrices (1× and 25×) in ln(1 + ΔF). W is a glomerulus × glomerulus matrix: diagonal entries relate a glomerulus to itself, off-diagonal entries to every other glomerulus. Estimating W entrywise would mean 141,376 coefficients from 59 odorants. We therefore fit it by partial least-squares regression (PLS-R, NIPALS; scikit-learn PLSRegression with scale=False), which summarizes the low-concentration population by a small number of latent activity patterns chosen to covary with the high-concentration response, then predicts every glomerular response from those patterns. Predictors and responses were mean-centered within each training fold without variance scaling; predictions were floored at zero. Eight components were selected by a one-standard-deviation rule on RMSECV over a 1–15 grid (the smallest number of components whose RMSECV lay within one standard deviation, taken across the grid, of the minimum); a one-standard-error rule and a Van der Voet randomization test (Van der Voet, 1994) both favored 11, and the structure of W was similar with 11 or 15 components.

All predicted responses were from held-out predictions: each odorant’s ‘high’-concentration responses were predicted from its ‘low’-concentration responses by the model fitted to the other three cross-validation folds. Cross-validation folds were stratified by chemical class, with classes of fewer than three odorants pooled. The weight matrix shown in **Fig. 7D**, fitted once to all 59 odorants, was used only to describe the overall learned transformation. The fitted model reached R²cv = 0.272, and 67% of glomeruli had positive individual R²cv. Permuting odorant labels of Y against X (1,000 permutations) gave a null of −0.226 ± 0.052, exceeded by every observed fit (p < 0.001).

For comparison with other regression methods, 18 alternative methods were run on identical data, folds, and preprocessing: ridge, RidgeCV, Lasso, elastic net, principal-component regression, reduced-rank regression, kernel ridge, random forest, extra trees, gradient boosting, multilayer perceptrons, k-nearest-neighbors, and identity and mean baselines. Four modestly exceeded the reported model in terms of variance explained (R²cv 0.276–0.343). We retained the low-rank linear fit using PLS-R because it yields an interpretable glomerulus × glomerulus operator.

#### Diagonal and non-diagonal contributions of W to prediction

W was partitioned into a block-diagonal region, comprising coefficients that link glomeruli within the same tuning cluster (22,404 of 141,376 entries), and a cross-cluster region. Coefficients outside a region were set to zero in each training fold’s fitted W and the masked operator applied to held-out odorants, so that all comparisons use the same fitted model. The block-diagonal region retained R²cv = 0.208, or 76% of the full model; cross-cluster coefficients alone retained 0.083, or 30%; and the literal diagonal retained essentially none (R²cv = −0.007). By contrast, predicting each glomerulus’s ‘high’-concentration responses from its own ‘low’-concentration responses alone, by a least-squares slope and intercept fitted to the training odorants of each fold, gave R²cv = 0.261.

Because a low-rank population makes any sufficiently large set of glomeruli near-equivalent as predictors, we compared the block-diagonal region with block-diagonal regions defined by randomly permuting cluster labels across glomeruli while preserving cluster sizes (200 permutations); these retained 26 ± 1% of the full model’s performance (p = 0.005), compared with 76% for the actual clusters.

#### Cluster-based contributions to low - to - high predictions

To assess the contribution of other canonical tuning motifs to predictions of concentration-dependent broadening (Fig. 8), the mean low-concentration profile of each of the seven clusters served as a reference (i.e., ‘source’) tuning motif, computed deterministically from X, so that no parameters are fit and no leakage occurs. For each of the 42 cluster pairs (source vs. non-source), every glomerulus of the source cluster was predicted by ridge regression from the source’s own motif alone, and from the source’s motif plus the non-source’s, under the same folds. The transfer statistic is the out-of-fold increment ΔR²cv = R²cv(source + non-source) − R²cv(source), pooled over the source cluster’s glomeruli. Because block structure in W can arise from correlations among low-concentration profiles alone, motif transfer was tested on held-out odorants.

The null distribution was built by shuffling the odorant ordering of Y against X (10,000 permutations), preserving all structure within X and within Y while destroying the low→high correspondence. One-sided p-values were computed as (1 + #{null ≥ observed}) / (1 + R) and corrected across the 42 pairs by the Benjamini–Hochberg procedure. Eight pairs were significant at FDR < 0.05.

#### Code and data availability

Raw data underlying all analyses, as well as a single code package used to generate all analyses and figures (generated by J. Castro and further modified by J. Hallkaj) can be found at https://github.com/WachowiakLab/OdorMapping2-Concentrations. Analysis code used Python 3.11 with scikit-learn 1.6.1, NumPy, SciPy, Matplotlib and Plotly. The PLS model, cross-validation, nulls, and regressor comparison are in github.com/CastroLab/pls-explorer; processed response data and cluster assignments are in github.com/CastroLab/glom-explorer. Random seeds are fixed and recorded in configs/baseline.yaml and in each analysis script.

## Acknowledgements

We thank Rebecca Kummer, Gustavo Opazo-Vasquez and Tom Rust for technical support.

---

**Table S1.**
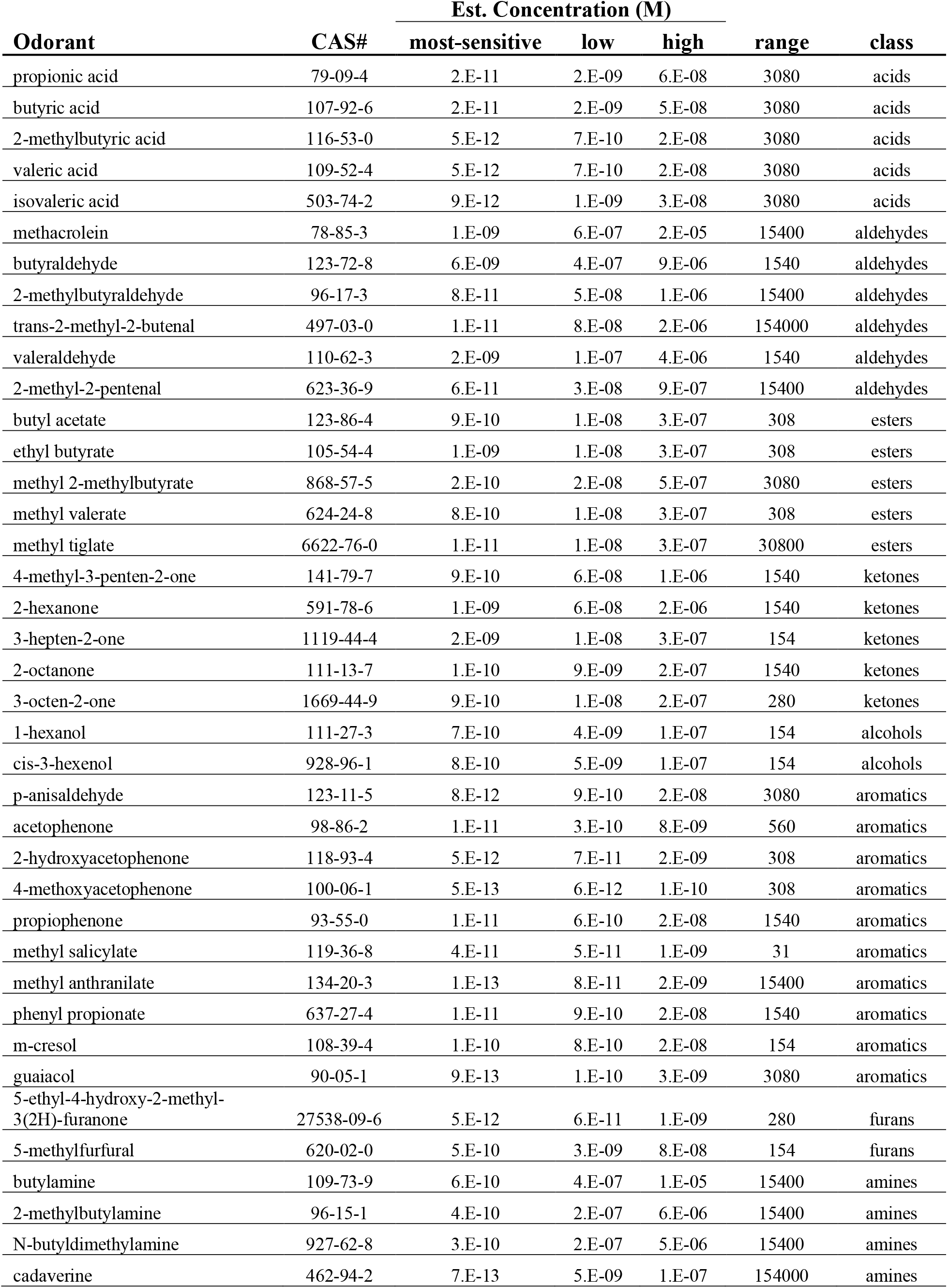
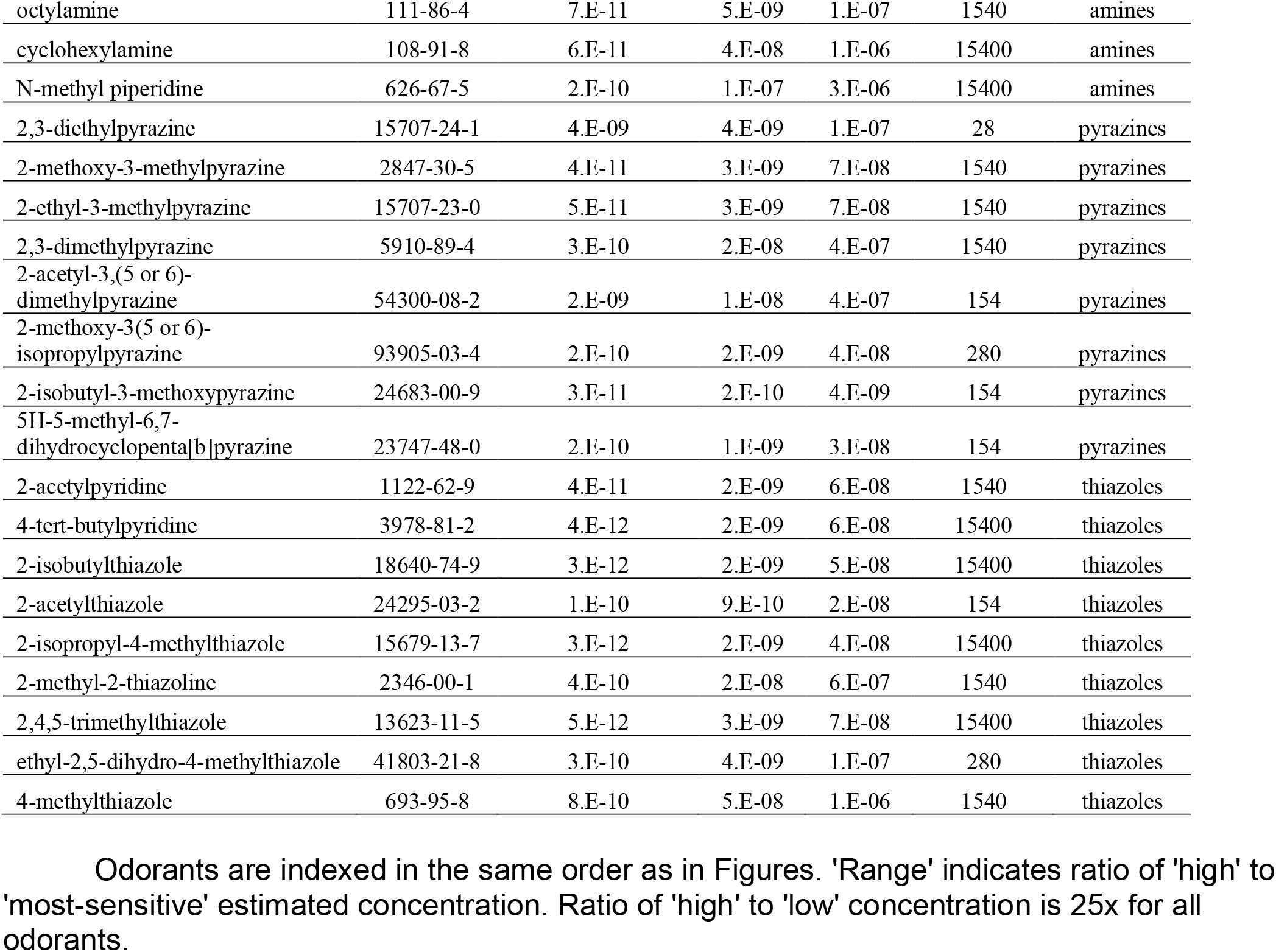
Odorants and estimated concentrations used in the study.

**Figure S1 (Related to Figure 4).**
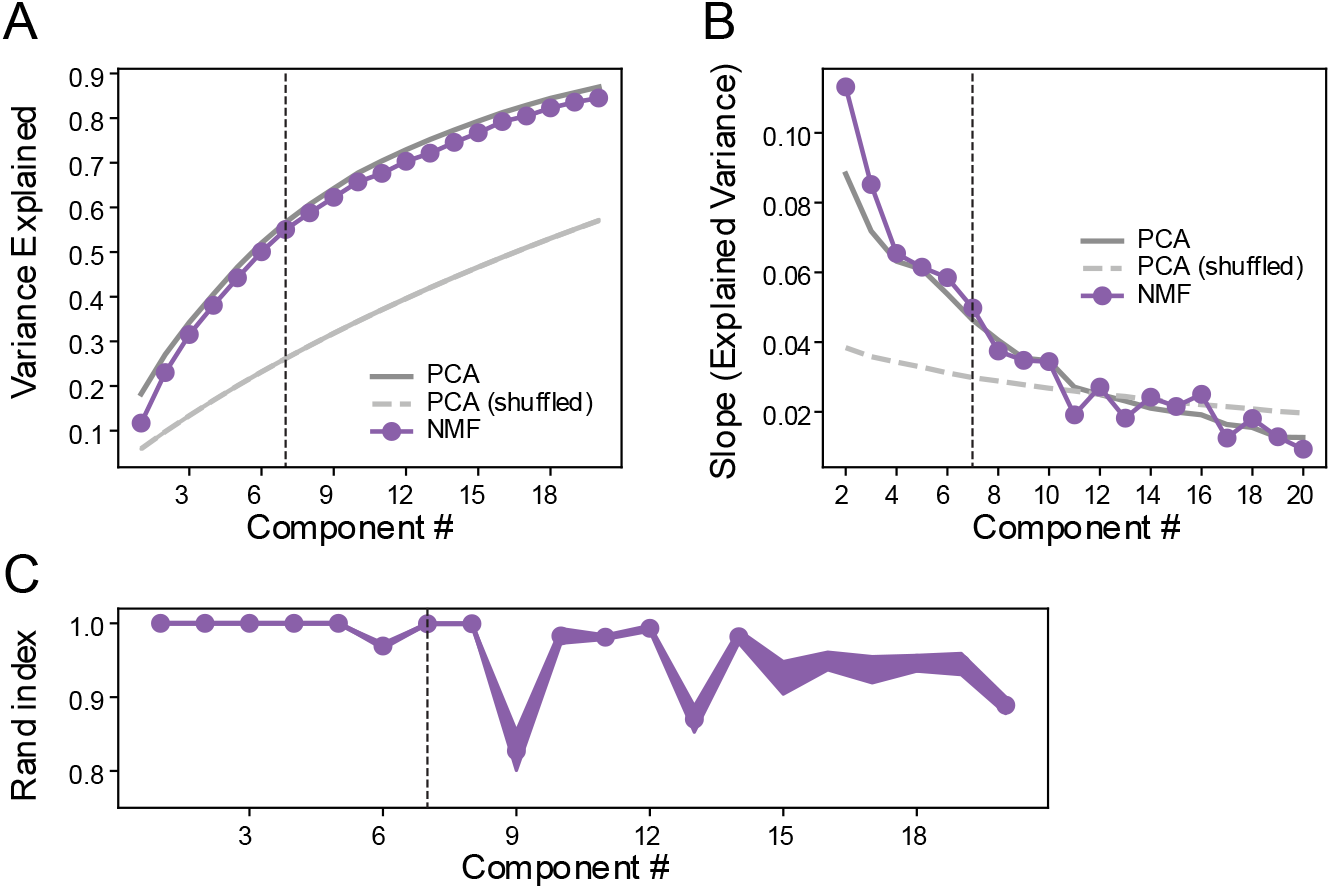
Choice of component number for NMF-based clustering of glomerular response vectors. **A.** Cumulative variance explained for dimensionality reduction of the ‘low’ concentration dataset pooled across subjects (59 odorants x 376 glomeruli). The gap between real and shuffled curves represents the structure not attributed to chance. **B.** Respective slope of the cumulative explained variance curves for the dimensionality reduction techniques, indicating how much additional components adds to the variance explained. **C.** Adjusted rand index across 50 different seeds of NMF runs, reflecting the stability of the clustering outcome.

**Figure S2 (Related to Figure 4).**
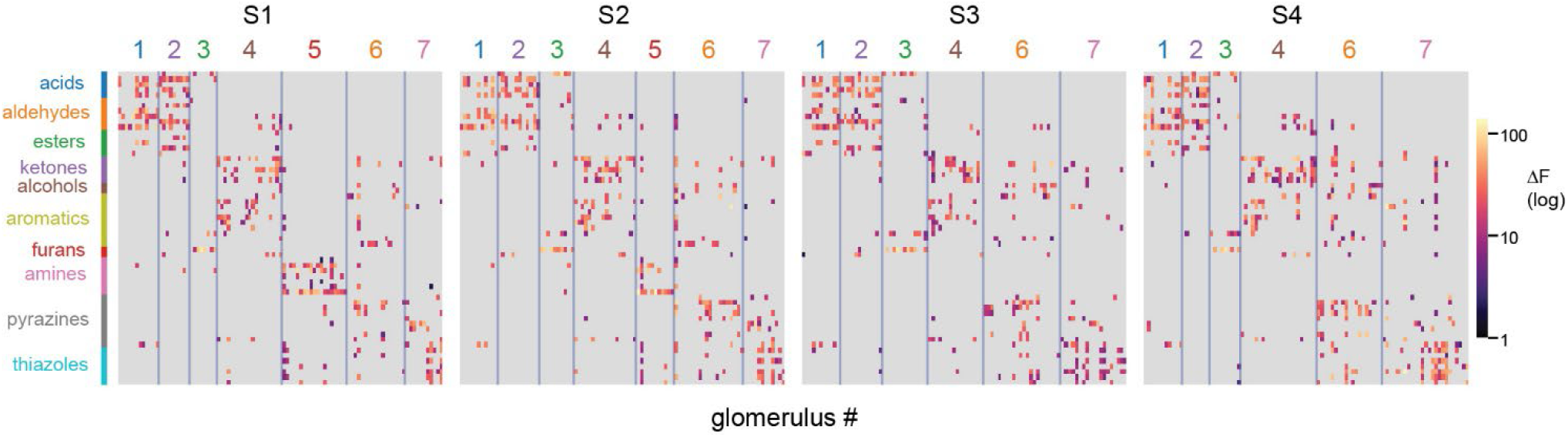
Odorant tuning structure of NMF clusters is preserved within each experimental subject. Odorant by glomerulus matrix of clustered responses, disaggregated by the four experimental subjects (S1 - S4). Blue lines represent NMF cluster borders across both glomerulus and odorant cluster. Panel depicts the same data shown in Figure 4A. Response amplitudes (ΔF) are mapped to a log pseudocolor scale for visualization.

**Figure S3 (Related to Figure 6).**
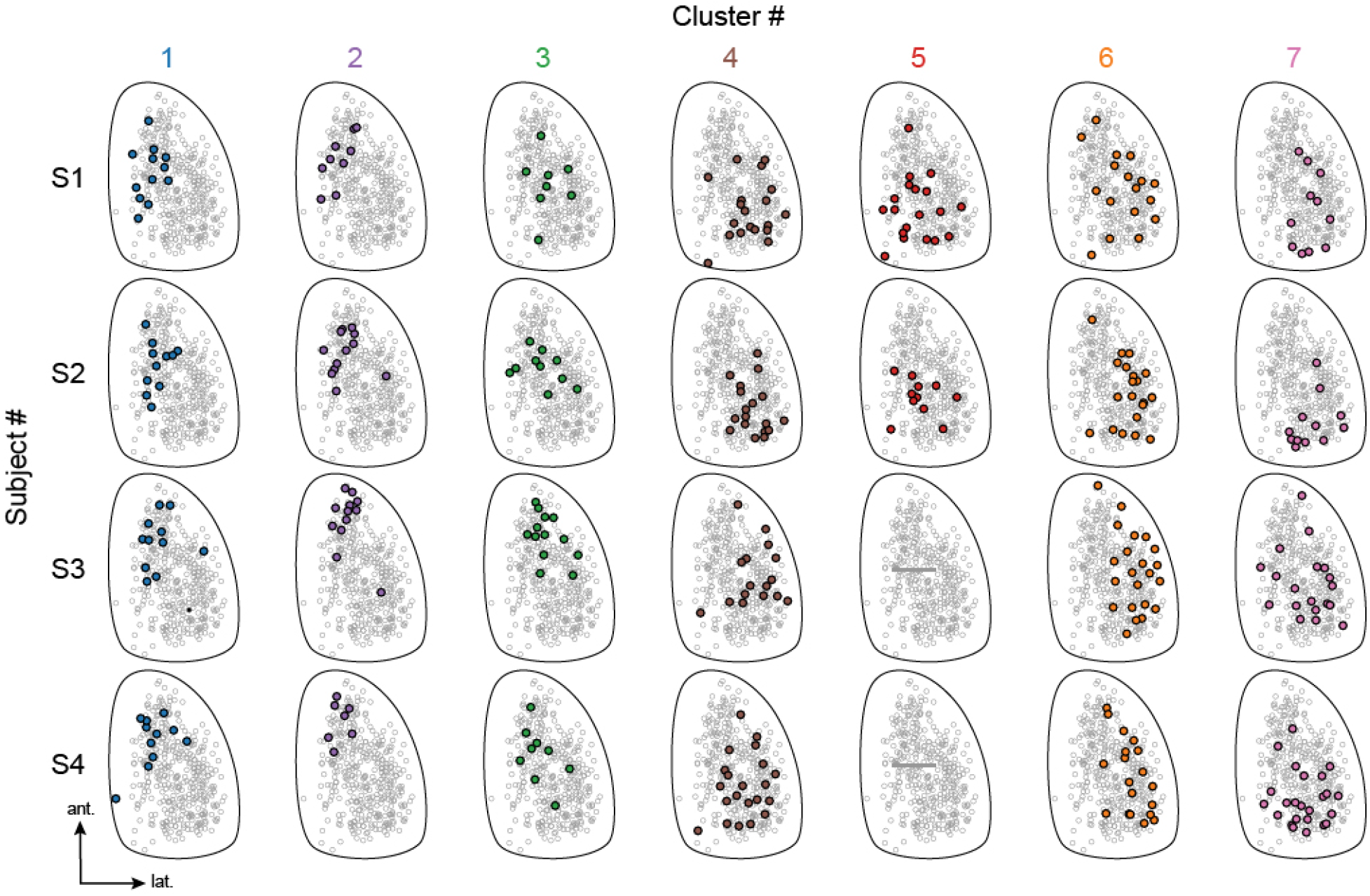
Locations of cluster-defined glomeruli are consistent in each OB. Locations of glomeruli belonging to each of the 7 clusters, disaggregated by experimental subject. Gray, unfilled circles represent all glomeruli pooled across subjects (same as in Fig. 6); filled, colored circles indicate responsive glomeruli from each subject (rows, S1 - S4). There were no glomeruli from cluster 5 in S3 and S4 as amines were not tested in these two subjects.

**Figure S4 (Related to Figure 7).**
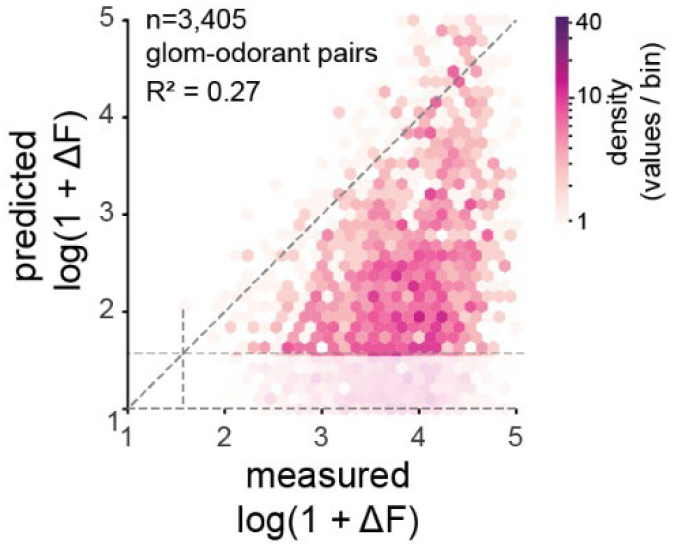
Measured vs. PLS-R-predicted high-concentration responses for all glomerulus-odorant pairs. Hexbin density plot of predicted versus measured responses for all individual glomerulus– odorant pairs with a non-zero response at high concentration (n = 3,405); pseudocolor scale indicates number of values within a given binned response range. Cross-validated R² over all pairs = 0.27. All predictions are held out (four-fold cross-validation over odorants). Predicted values below the minimum measured response amplitude are rendered partially transparent.

